# Nitrogen source and Nod factor signaling map out the assemblies of *Lotus japonicus* root bacterial communities

**DOI:** 10.1101/2023.05.27.542319

**Authors:** Ke Tao, Ib T. Jensen, Sha Zhang, Eber Villa-Rodríguez, Zuzana Blahovska, Camilla Lind Salomonsen, Anna Martyn, Þuríður Nótt Björgvinsdóttir, Simon Kelly, Luc Janss, Marianne Glasius, Rasmus Waagepetersen, Simona Radutoiu

**Affiliations:** Department of Molecular Biology and Genetics, Aarhus University, Denmark; Department of Chemistry, Aarhus University, Denmark; Center for Quantitative Genetics and Genomics, Aarhus University, Denmark; Department of Mathematical Sciences, Aalborg University, Denmark

## Abstract

Symbiosis with soil-dwelling bacteria that fix atmospheric nitrogen allows legume plants to grow in nitrogen-depleted soil. Symbiosis impacts the assembly of root microbiota, but it is not known how this process takes place and whether it is independent of nitrogen nutrition. We use plant and bacterial mutants to address the role of Nod factor signaling on *Lotus japonicus* root microbiota assembly. We found that Nod factors are produced by symbionts to activate Nod factor signaling in the host, and this modulates the assembly of a symbiotic root microbiota. *Lotus* plants grown in symbiosis-permissive or suppressive soils delineated three nitrogen-dependent nutritional states: starved, symbiotic, or inorganic. We found that root and rhizosphere microbiomes associated with these states differ in composition and connectivity, demonstrating that symbiosis and inorganic nitrogen impact the legume root microbiota differently. Finally, we demonstrated that selected bacterial genera delineating state-dependent microbiomes have a high level of accurate prediction.

## Introduction

Plant tissues are habitats for communities of microbes, generally called the plant microbiota [1–3]. Members of these communities may colonize the intra- or intercellular space of the plants, as well as the surface of aerial and belowground organs [3–6]. Microbial colonization of plant tissues involves an extensive selection, and evidence for an active role of the host in this process has emerged from studies across numerous plant species and growth conditions [7–11]. Phylogenetically distinct plant species host different microbial communities, which primarily differentiate at a low taxonomic level [10, 12, 13]. Reconstitution experiments using synthetic communities revealed that bacterial isolates assigned to the same family and originating from the same soil may have preferences for colonizing the roots of a legume or brassica [12]. This host preference was observed only in the presence of metabolically active roots suggesting microbial adaptation to plant-derived environmental niches. Plant species differ in root morphology, cell wall composition as well as the pattern and magnitude of metabolites secreted in the rhizosphere, which in turn impact the associated microbiota [14–19]. These traits also vary within the same species depending on plant genotype or physiological state. The nutritional state of the host was consistently identified to impact microbiota composition across plant species [20, 21]. In nutrient deplete state, plants adjust the profile of secreted metabolites which leads to the enrichment of specific members from soil microbiota [22–24], some engaging in symbiotic associations with the host [25–27].

Legume plants are genetically equipped to enrich their rhizosphere with soil bacteria that fix atmospheric nitrogen which are accommodated inside their roots and root nodules [25, 27, 28] and can thus overcome the lack of nitrogen in the soil. Most of these symbionts do not fix nitrogen as free-living bacteria in the soil, and legume plants select them based on complex signal exchange and recognition [29–31]. Consequently, bacterial mutants impaired in nitrogen fixation but able to produce compatible symbiotic signals, the Nod factors, can colonize legume roots and nodules efficiently [32, 33]. Host-specific flavonoids are released from roots into the soil and, if recognized by symbionts, induce expression of bacterial genes for Nod factor biosynthesis [34, 35]. Nod factors with species-specific decorations are recognized by plant receptors localized on the plasma membrane [30, 31, 33, 36]. The NFR1 and NFR5 receptors initiate symbiosis signaling in *Lotus japonicus* roots and initiate nodule organogenesis and infection thread formation [29, 37]. Symbionts infect the roots and nodules via infection threads [5, 38] where they continue to produce Nod factors while the infection threads progress inwards, across plant root cell layers and inside nodule primordia [28, 39]. The epidermal Nod factor receptor NFRe aids the symbiosis initiated by NFR1 and NFR5, enabling optimal signaling during nodulation and infection [40], while the Chitinase *Chit5* cleaves bacterial Nod factors, maintaining an optimal level of signaling during root and nodule primordia infection [41].

Previous studies revealed that the host pathway required for symbiosis with nitrogen-fixing bacteria has a major impact on root microbiota assembly of legume plants [7, 42–45], an effect which was retained in the presence of nitrogen-replete soil [7]. However, how symbiosis signaling, and nitrogen nutrition affect legume root microbiota structure remains unknown. Root-nodule symbiosis is an energy-demanding process for the host, and it is thus inhibited in nitrogen-replete conditions [46]. Therefore, when considering nitrogen nutrition, legumes can be found in three distinct states: starved, replete due to nitrogen-fixing symbiosis, or replete due to uptake of inorganic nitrogen from the soil. Both nitrogen-replete states ensure efficient plant growth and seed production, but legumes differ in the amount of nitrogen they require for high seed yield [47, 48]. Legumes and their symbionts thus provide an appropriate system for investigating if acquisition of nitrogen through symbiosis with a soil microbe or directly from the soil reserves have an impact on microbiota assembly and how symbiosis signaling, and nitrogen source contribute to root microbiota establishment.

Here, we use *Lotus* and *Mesorhizobium loti* mutants to address these questions. Our experiments using soil permissive or suppressive for symbiosis, as well as those based on gnotobiotic settings with synthetic communities identified that nitrogen nutrition and Nod factor signaling are major drivers of *Lotus* root microbiota. We found that root microbiota is dependent on nitrogen-nutrition and source and that bacterial genera characterizing these states have high prediction power. Importantly, our results based on Nod factor signaling in *Lotus* provide evidence that interactions established between distinct members of soil microbiota and the host can lead to a feedback effect on the remaining members of the community which is orchestrated by the host.

## Results

### The growth of *Lotus* in soil is supported by inorganic nitrogen and symbiosis

Nitrogen-fixing symbiosis is a facultative trait. Legumes initiate symbiotic signaling and allow bacteria to infect their roots if nitrogen in the soil is insufficient for plant growth. To investigate if acquisition of nitrogen through symbiosis or from the soil reserves has a differential impact on the assembly of bacterial communities of *Lotus,* we exposed wild-type plants (Gifu) and three mutants impaired in symbiosis signaling to growth in native (symbiosis permissive) or nitrate-supplemented (symbiosis-inhibiting via10 mM KNO_3_) agricultural Cologne soil, under controlled conditions (Fig. 1a). Plants were evaluated for growth and symbiotic phenotype in the presence of native soil microbiota after nine weeks of growth. The three mutants are impaired in Nod factor signaling acting at different stages of symbiosis [29, 40, 41]. Mutation in the *Nfr5* impairs Nod factor perception and initiation of all symbiotic events in the root [29, 37]. Mutants in *Nfre* are impaired in Nod factor perception in the epidermis and have a suboptimal symbiotic phenotype. Functional nitrogen-fixing nodules are formed, albeit in a reduced number compared to Gifu [40]. The *chit5* mutants lack the ability to balance the levels of Nod factor signaling during cortical infection. Nodules are formed, but most of them are non- or poorly infected [41]. We observed that mutants grown in natural soil showed reduced shoot biomass, and only Gifu and *nfre* reached the reproductive stage. (Fig. 1b). This reduction in shoot biomass was likely a result of the gradual decrease in the number of functional nodules, leading to nitrogen-starved *nfr5* and *chit5* plants (Fig. 1d and 1e). On the other hand, nitrate supplementation of the soil ensured flowering and high shoot biomass for all genotypes (Fig. 1c), indicating plentiful nitrogen nutrition independent of symbiosis impairment. In this condition, Gifu, *nfre*, and *chit5* plants formed a reduced number of sparsely distributed white nodules (Fig. 1f). No functional pink nodules were detected, reflecting a dramatically reduced symbiosis.

**Figure 1.**
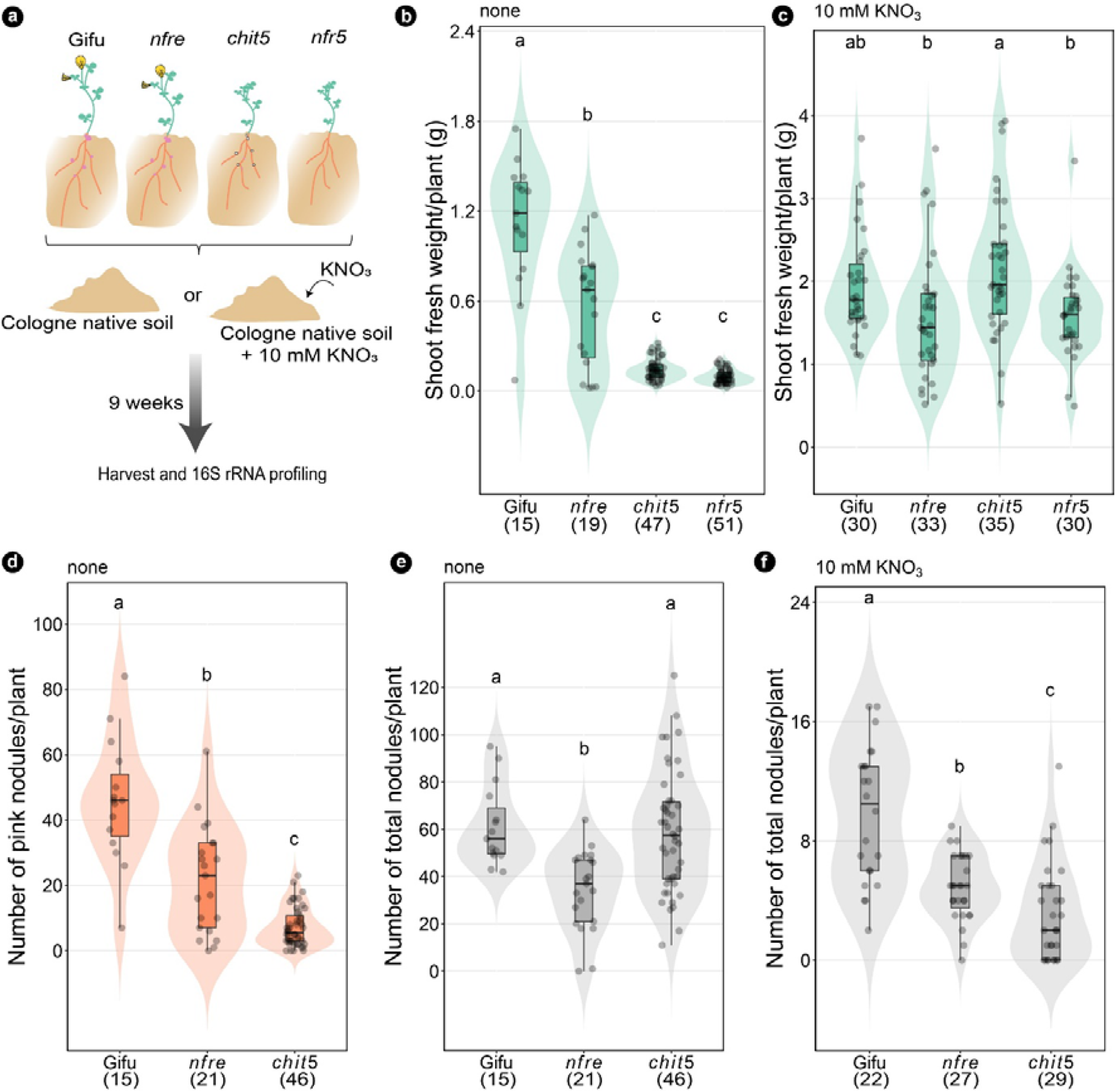
Wild-type and Nod factor signaling mutants grown in soil show symbiosis defective phenotypes compensated by nitrate addition to the soil. (**a**). Experimental design. (**b**) Shoot fresh weight, the number of pink nodules (**d**), and the total number of nodules (**e**) per plant of wild-type, *nfre*, *chit5*, and *nfr5* grown in native Cologne soil. Shoot fresh weight (**c**), and total number of nodules (**f**) of wild-type, *nfre, chit5,* and *nfr5* grown in native Cologne soil supplied with 10 mM KNO_3_. Each dot represents values for individual plants. The shape of violins illustrates the density of samples. Boxplots within the violin plots show the median, 75^th^ percentile, and 25^th^ percentile datasets. Letters within the plots (**b, c, d, e, f**) indicate statistically significant differences (Tukey HSD test, *p*<0.05). The number of plants analyzed for each genotype is in brackets. Note the differences in the Y axes for panels (**e**) and (**f**).

These phenotypes of soil-grown plants show that both nitrate and the efficient nitrogen-fixing symbiosis are conducive conditions for healthy and reproductive *Lotus* plants, and that NFRe, CHIT5, and NFR5 proteins contribute differently to root nodule symbiosis and nitrogen nutrition.

### The physiological state of *Lotus* has an impact on the associated bacterial communities

Next, we used 16S rRNA amplicon sequences, to investigate the structure of bacterial communities assembled in the root and rhizosphere compartments of Gifu and symbiotic mutants grown in native or nitrate-supplemented soil. The addition of inorganic nitrogen reduced the α-diversity of the unplanted soil, as well as in the rhizosphere of *nfr5* and *chit5*, while the remaining samples maintained a similar diversity within the sample in the presence of nitrate (Fig. 2a and Supplementary Fig. 1a). The β-diversity analysis based on Bray-Curtis dissimilarities (between sample diversity), showed a clear separation of communities present in the rhizosphere and root samples (Supplementary Fig. 1b). Overall, the interactions between nitrate supplementation and the genotype account for 44.8% and 36.2% of the variation in the root and rhizosphere compartments respectively (Fig. 2b and 2c). The nitrate had a large impact on both root and rhizosphere communities with a stronger effect on the rhizosphere (R^2^=0.297 versus R^2^=0.220 in the root), while the genotype had a larger impact on the root communities (R^2^=0.140 versus R^2^=0.119 in the rhizosphere) (Supplementary Fig. 1e and 1f). The principal coordinate analysis (PCoA) with Bray-Curtis distances for rhizosphere samples showed a clear separation of the analyzed communities into three clusters. One cluster contained communities of Gifu and *nfre* plants grown in natural soil supported by nitrogen-fixing symbiosis. The second cluster contained communities of nitrogen-starved plants: *chit5* and *nfr5* grown in natural soil. The third cluster contained communities of all genotypes grown in soil supplemented with nitrate (Fig. 2b). A similar separation could be identified for communities present in the root samples, (Fig. 2c). These differences in bacterial communities followed the observed phenotypes (Fig. 1b and 1c) and the physiological state of the host—nitrogen starved, symbiotically active, and inorganic nitrogen-supported plants.

**Figure 2.**
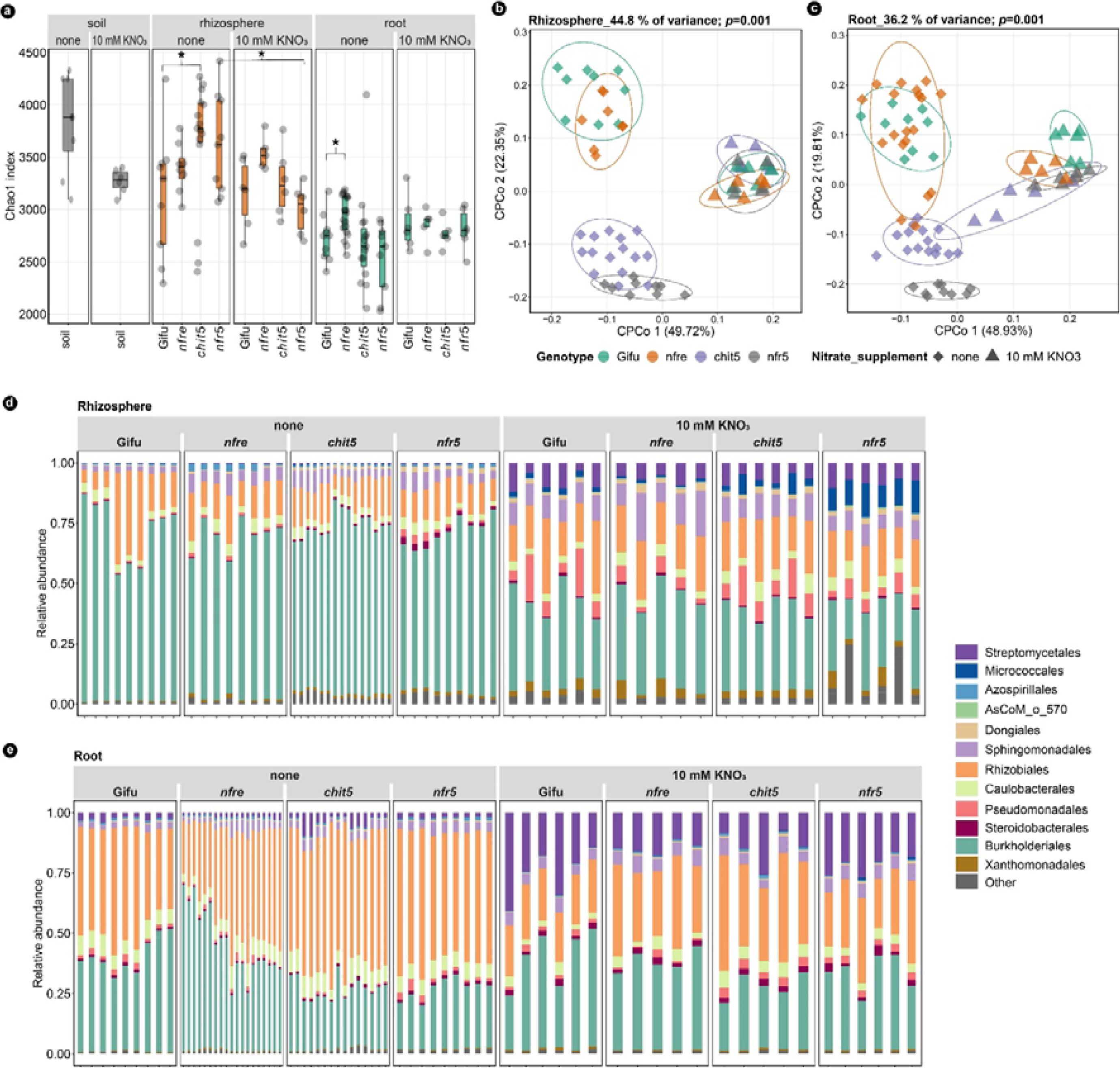
Nitrate supplementation changes the soil, root, and rhizosphere community structures. **(a)** Alpha diversity by Chao1 index for soil, rhizosphere, and root compartments of Gifu, *nfre*, *chit5*, and *nfr5.* (**b)** Constrained PCoAs of rhizosphere and root (**c**) communities show that Nod factor signaling, and nitrate supplementation have a major effect on the associated bacteria. The analysis was constrained by both genotype and nitrate application. (**d**) Relative abundance of bacterial families for rhizosphere and root (**e**) samples. Column indicates the replica and colors indicate the taxonomic assignment.

Next, we analyzed if differences observed at the community level associate with changes in the relative abundance (RA) of specific taxa. The 16S rRNA amplicons were clustered into ASVs that were taxonomically assigned using the ecosystem specific database AsCoM [49], and corresponding RAs were calculated. Comparative analyses identified significant shifts in the RA of Burkholderiales, Rhizobiales, and Streptomycetales (Fig. 2d and 2e, Supplementary Fig 2) in samples grown on native versus nitrate-supplemented soil, indicating that the type of nitrogen nutrition impacts the associated communities at high taxonomic level. By contrast, no obvious differences were identified between Gifu and symbiotic mutants grown in the native soil when comparing the RA of different bacterial orders (Fig. 2d, 2e and Supplementary Fig. 2c, 2d).

Together, these results from analyses of α- and β-diversity, as well as from comparing differential abundances of bacterial orders show that nutritional state (nitrogen deplete or replete) as well as the source of nitrogen (symbiotic or inorganic) are major factors differentiating bacterial communities in soil-grown *Lotus* plants. Importantly, bacterial communities associated with symbiotically active plants differ from those associated with plants supplied with inorganic nitrogen, even if plants are in a nitrogen-replete state in both conditions.

### Mutations in Nod factor signaling have a differential impact on root microbiomes

Diversity analyses revealed a clear separation of communities associated with starved (*nfr5* and *chit5*) versus symbiotic (Gifu and *nfre*) plants, but no significant differences were detected at the order level (Fig. 2e). Thus, we set out to investigate if significant differences are detected between root communities of wild-type and symbiotic mutants grown in native Cologne soil at lower taxonomic levels. We focused primarily on ASVs with RA larger than 0.3% (n=141) in Gifu roots (Fig. 3). Together, these have a cumulative RA larger than 60% (Fig. 3b) and are assigned to 10 bacterial orders and 14 families, with members of Rhizobiales, Caulobacterales, and Burkholderiales in majority (Fig. 3b and Supplementary Table 1). Communities of *nfre* were found to be most similar to Gifu, and only 11 ASVs were significantly reduced (Fig. 3d and Supplementary Table 1). By contrast, 31 ASVs had a significantly reduced abundance in *chit5*, and 48 ASVs in the *nfr5* roots. Most importantly, these ASVs belong to families that are highly abundant in Gifu roots. Several ASVs assigned to *Oxalobacteraceae*, *Comamonadaceae*, *Rhizobiaceae*, and *Xanthobacteraceae* were significantly reduced in abundance in all mutants (Supplementary Table 1). Moreover, the RA of *Mesorhizobium*, *Oxalobacteraceae*, *AsCoM*_f_4827 (Gammaproteobacteria), together accounting for more than 15% abundance in wild-type, were significantly reduced in *chit5* and *nfr5* (Fig. 3c and 3d, Supplementary Fig. 4, Supplementary Table 2).

**Figure 3.**
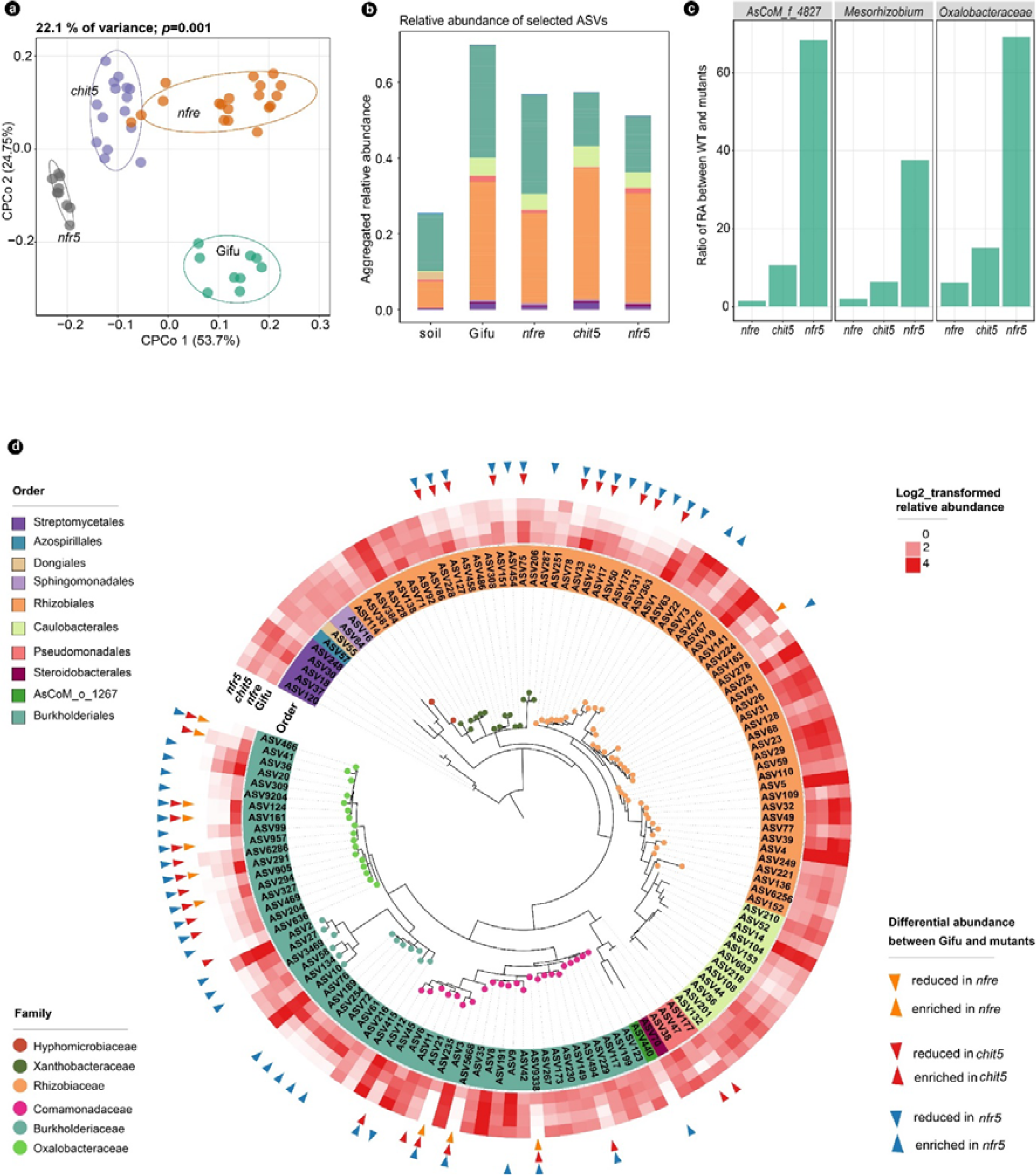
Nod factor signaling contributes to root-associated microbiota of *Lotus* plants grown in native Cologne soil. **a**) Constrained PCoA of communities associated with roots of wild-type, *nfr5*, *nfre* and *chit5*. **b)** Cumulative relative abundance of selected ASVs (RA >0.3% in roots of Gifu) in soil and roots of the four genotypes. **c**) The ratio of RA between Gifu and mutants of the top three taxa in the roots based on selected ASVs: RA>0.3% in roots of Gifu. **d**) Distinct ASVs have a significantly different RA in mutant roots compared to wild-type Gifu. Selected ASVs are presented in a phylogenetic tree constructed on the basis of 16S rRNA V5-V7 region. The taxonomic information is shown by color on the name of the ASV (order) and by color on the tree branch (family). The heatmap shows the log2- transformed RA of each ASVs in the roots of Gifu, *nfre*, *chit5*, and *nfr5* plants. Triangles on the outer layer of the heatmap point out ASVs that have a significantly different abundance compared to wild-type plants.

This shows that a gradual impairment of Nod factor signaling has a corresponding impact on root microbiota and that host genotype affects the composition of bacterial communities present in *Lotus* roots grown in symbiotic permissive soil, as indicated by PERMANOVA analyses (Supplementary Fig. 1f).

### Supplementation of soil with inorganic nitrogen leads to changes in root and rhizosphere microbiomes

Our analyses of community structures identified a significant effect of soil supplementation with inorganic nitrogen (Supplementary Fig. 6–9). We analyzed these changes at the composition level and found that nitrate application had an impact on the unplanted soil microbiota, where 39 ASVs assigned to 13 families had a reduced abundance compared to the native soil (Supplementary Fig. 6–9, Supplementary Table 3) explaining the observed reduction in α-diversity (Fig. 2a and Supplementary Fig. 1a). This shows that nitrate application has a direct impact on some of the soil bacteria (Supplementary Table 3). Surprisingly, we found a much larger impact of nitrate application onto the rhizosphere and root communities of all plant genotypes, and that ASVs affected in the unplanted soil represented only a minor fraction of the overall changes detected in the root and rhizosphere compartments (Supplementary Fig. 6–9). We found that the order Burkholderiales which dominated the rhizosphere communities of plants grown in native soil, was largely affected and substantially reduced in abundance when plants were grown in nitrate-supplemented condition (Fig. 2d and Supplementary Fig. 2c). This reduction was balanced by an increase in abundance of three bacterial orders, Pseudomonadales, Streptomycetales, and Sphingomondales (Fig. 2d and Supplementary Fig. 2c). In the root compartments, we found that Rhizobiales which was the most abundant taxon when plants were grown in natural soil (Fig. 2e and Supplementary Fig. 2d) was substantially reduced. This reduction was compensated for by an increase in the abundance of Streptomycetales.

In addition to these changes in the cumulative abundances of major bacterial orders, plant growth on inorganic nitrogen had a clear effect on the abundance of individual ASVs. Consequently, the composition of ASVs within the same families changed, indicating a possible shift in the functional capacities at the family level (Supplementary Fig. 6–9).

Bacterial communities of wild-type and mutants grown in nitrate-supplemented soil clustered together, indicating a minor effect of the genotype in these conditions (Fig 2b and 2c). Indeed, when comparing wild-type and mutants we found several ASVs having a different abundance, but these were not statistically significant (Supplementary Fig. 3). Only when analyzing the cumulative abundance of ASVs belonging to *Mesorhizobium*, *Burkholderiaceae*, and *Nocardioiaceae* families could we detect a significant reduction in at least one of the symbiotic mutants (Supplementary Fig. 3c and Supplementary Fig. 5).

Together, these results show that supplementation of soils with inorganic nitrogen has a consequential impact on the community structures and composition in the rhizosphere and root of *Lotus*, and importantly, these communities differ significantly from those associated with nitrogen-fixing plants.

### Plants grown in symbiosis suppressive conditions have microbiomes with reduced connectivity

Growth of wild-type and mutants in symbiosis-permissive or suppressive soil identified three types of microbiome assemblies in both roots and rhizosphere compartments. (Fig. 2b and 2c). This indicates that nitrogen nutrition of *Lotus* plants delineates distinct environmental niches that may feature specific microbial co-occurrence networks and relationships. To study these, we analyzed the co-occurrence patterns for bacterial ASVs identified in the rhizosphere and root compartments of plants, defining the three states (Fig. 2b and 2c) (see methods). The starved state was defined by communities of *nfr5* and *chit5* grown in native soil, the symbiotic state by those of Gifu and *nfre* plants grown in native soil, while the inorganic nitrogen state was defined by communities of all genotypes grown in nitrate-supplemented condition.

We found that networks characterizing communities of plants grown in nitrate-supplemented soil had a lower number of edges compared to those of symbiotic or starved plants, and that nodes of the inferred root and rhizosphere networks were primarily isolated (Fig. 4a and 4b, Supplementary Fig. 10c). The modules identified in the networks for rhizosphere (n=3) and root (n=6) communities contained a limited number of vertices (Supplementary Table 4), which in general belonged to a dominating genus (Fig. 4a1 and 4b1). This indicates that communities associated with *Lotus* grown on inorganic nitrogen are composed of isolated nodes that establish few connections with each other, and most of the established connections are between nodes belonging primarily to the same genus.

**Figure 4.**
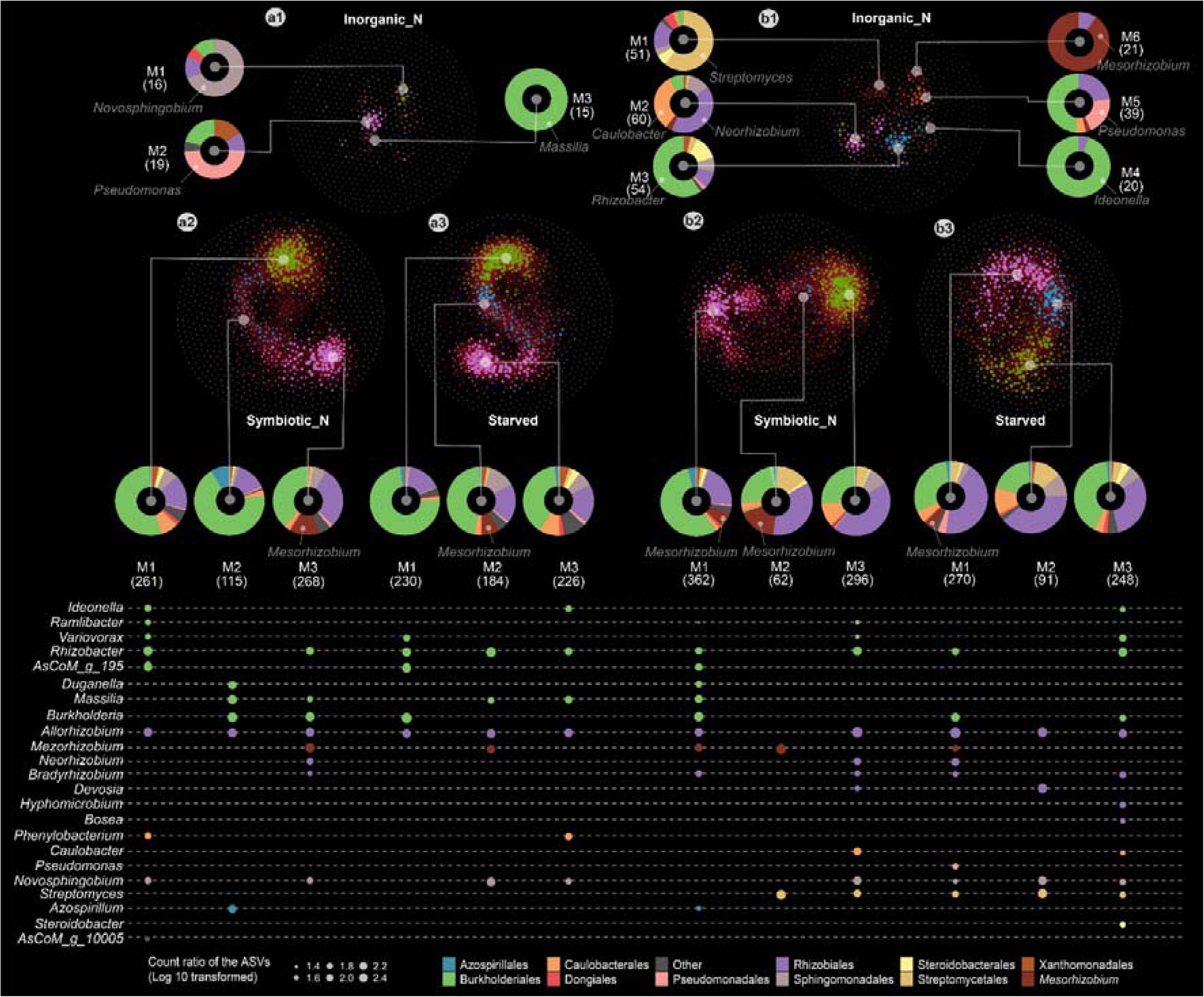
Nitrogen nutritional status and source drive distinct correlation networks between ASVs into the rhizosphere and roots of *Lotus.* Correlation networks of ASVs in the rhizosphere (**a**) and root (**b**) compartments of plants supplemented with inorganic nitrogen (**a1** and **b1**), or symbiotic nitrogen (**a2** and **b2**) source, or starved (**a3** and **b3**) status. Only positive correlations are marked out by red lines between nodes on the networks. The nodes of the networks are colored by modularity class. Each node represents an ASV, the size of the node is given by its degree (the degree of a node refers to the number of other nodes it is connected to). The modularity classes are denoted M1, M2, M3, M4, M5, and M6, and the numbers of ASVs in each module are shown in brackets. For each module, the proportion of ASVs at the taxonomic order level is shown by donut plots. The main taxonomic genus in each modularity is pointed out by text on the donuts (**a1** and **b1**) or dots below the modularity (**a2**, **a3**, **b2**, **b3**). The color of the dots represents the taxonomic order. The size of the dots represents the percentage of ASVs in the module.

The networks identified for communities from rhizosphere and roots of symbiotic (Gifu and *nfre*) or starved plants (*nfr5* and *chit5*) contained a larger number of edges and more than 60% of nodes were found nested into a module (Fig. 4a2, 4a3, 4b2 and 4b3, Supplementary Fig. 10). The positive edges established between nodes drove the assemblies into three major modules (Fig. 4a2, 4a3, 4b2 and 4b3), while negative edges were primarily established between members of different modules (Supplementary Table 4). Bacterial ASVs belonging to Burkholderiales and Rhizobiales were identified as hubs in all major modules, and interestingly, the ASV1 which is most likely the symbiont (Supplementary Fig. 10) was not among them (Supplementary Table 4). The composition and taxonomic representation of ASVs within modules varied, indicating that different interactions may establish between ASVs in the root and rhizosphere (Fig. 4a, 4b and Supplementary Fig. 10). As an example, we found that ASVs belonging to eight genera of Burkholderiales assembled differently in the root and rhizosphere modules (Fig. 4a2 and 4a3). By contrast, the Rhizobiales order dominated two (M2, M3) of the three major modules identified in the root communities.

Symbiotic plants (Gifu and *nfre*) contained numerous ASVs belonging to Burkholderia genus (n=27) in the modules identified in both rhizosphere and root. These ASVs were absent in the corresponding modules of starved plants, where the *Rhizobacter* genus was found instead (n=6). The *Mesorhizobium* genus was largely represented in the root and rhizosphere modules of symbiotic plants (n=26, of which one is the symbiont), but its representation was reduced in the corresponding modules of the starved plants (n=6), where a larger number of ASVs assigned to *Allorhizobium* (n=12) were identified. This indicates that functional symbiosis through Nod factor signaling promotes an environmental niche where distinct members of *Mesorhizobium* and *Burkholderia* genera may engage in positive interactions with the symbiont. This finding prompted us to ask if such inferred positive interactions were established, which would lead to the expectation the these ASVs are affected in their abundance by the impairment of Nod factor signaling in the mutants. Indeed, we found that many of the ASVs present in the same module as ASV1 (rhizosphere M3 and root M1) of symbiotic plants, had a lower abundance in the *nfr5* and *chit5* mutants (Fig. 3 and Supplementary Table 1, 4).

Together, the co-occurrence networks revealed that bacteria are highly interconnected in communities associated with symbiotic and starved plants, while those of plants grown on inorganic nitrogen establish fewer and different interactions. Furthermore, this analysis identified members of microbiota that engage in positive interactions with the symbiont if conditions are conducive for an optimal Nod factor signaling and symbiosis.

### Specific genera can predict the nitrogen nutritional status of *Lotus* with high accuracy

The observed large differences in composition and connectivity between communities associated with *Lotus* plants supported by different nitrogen nutrition mode prompted us to determine if these are part of a more general, possibly predictable pattern. First, we identified significantly enriched and depleted taxa associated with the three states and evaluated their potential use for predicting the nitrogen-dependent state in unrelated/unknown microbiome of the same host grown in the same soil and conditions (Fig. 5). We conducted ternary comparisons between communities associated with (1) all plants grown on soil with inorganic nitrogen, (2) symbiotically active plants Gifu and *nfre* grown in native soil, and (3) nitrogen starved *chit5* and *nfr5* plants grown in native soil for both rhizosphere and root compartments This identified ASVs specifically enriched in the three conditions (Fig. 5a and 5b). Next, we aggregated the ASVs at the genus level (Supplementary Table 5) and identified 51 and 43 genera in the rhizosphere and root, respectively, enriched or depleted in at least one of the three nutritional states (Fig. 5c and 5d, Supplementary Table 5). These were used as covariates to predict the nitrogen nutritional state of *Lotus* plants from an independent, but similar dataset described by Zgadzaj et al. [7]. Samples from Zgadzaj et al. were split into a training and test dataset (Fig. 5c and 5d). A filtering step was applied separately to training and test data, followed by centered log ratio transformation of the relative abundances of each genus, and a variable selection procedure [50] before the final model fitting (Fig. 5c and methods). Of the initial 51 significant genera identified in the rhizosphere dataset of this study, 28 were retained after variable selection (Fig. 5c and Supplementary Table 5). These were used as covariates in several prediction models fitted on the training data (Supplementary Table 5). The accuracy of the fitted models was assessed on the test dataset, and the best performance was obtained from a support vector machine with an accuracy of 79% (Fig. 5c). The same analysis was repeated on the root samples (Fig. 5d). Of the initial 43 genera, 23 were retained after variable selection (Supplementary Table 5), and here the support vector machine yielded a 74% accuracy on the root test dataset (Supplementary Table 5). These results indicate that genera identified to be enriched in one of the three states presented in this study have a high level of prediction of the nitrogen nutritional state of *Lotus* plants.

**Figure 5.**
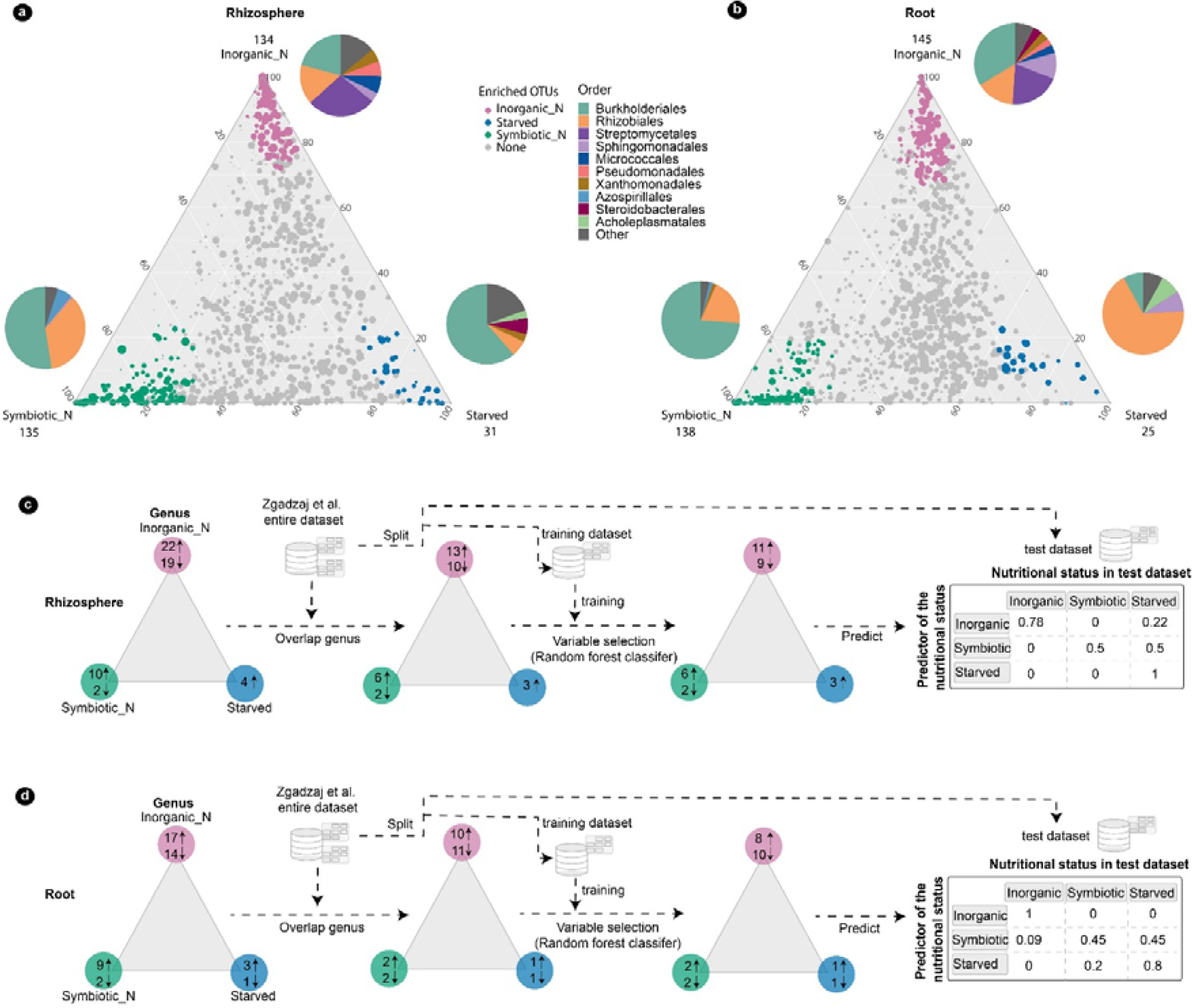
Specific bacterial taxa enriched in the three nutritional statuses are identified as predictors with high accuracy. Ternary plots illustrating the RA of all ASVs identified in the rhizosphere **(a)** or root (**b**) compartments of nitrogen-replete (inorganic nitrate- KNO3 or symbiotic nitrogen) or -deplete (starved) samples. Starved conditions are represented by the *nfr5* and *chit5* grown in native Cologne soil, symbiotic nitrogen conditions are represented by the wild-type and *nfre* grown in native Cologne soil, and inorganic nitrogen conditions are represented by all genotypes grown in nitrate supplemented Cologne soil. The ASVs are presented by dots; the size of the dots is determined by the mean RA across all three conditions in the ternary plots. The position of the dots in the ternary plot is determined by the mean RA of the ASVs within the three states. Green ASVs are enriched in symbiotic nitrogen condition, blue in starved condition, and pink in inorganic nitrogen condition. The number of enriched ASVs are marked at the corners. Pie charts show the order-level taxonomic composition of the enriched ASVs. **(c)** Scheme of the process to identify predictor taxa in the rhizosphere and **(d)** root. The triangle in the scheme is a simplified symbol of the ternary plot in **(a)** and **(b)**. Pink indicates inorganic nitrogen conditions, green indicates symbiotic nitrogen conditions, and blue indicates nitrogen starved conditions. Numbers in the circles indicate the number of enriched (upwards) or depleted (downwards) bacterial genera.

### Microbiota of *Lotus* is shaped by the presence of a Nod factor-producing symbiont

Results from current and previous studies [7, 43, 45] revealed that in the absence of a functional root nodule symbiosis, *Lotus* plants grown in native soil are nitrogen starved and assemble a distinct microbiota. However, it is unknown if nitrogen nutrition provided by the symbiont is the sole determinant for the observed differences. Controlling the presence of the symbiont or its functional capacities at the native soil level is not feasible, thus, to answer this question we performed a series of studies using synthetic communities. We assembled a “symbiont-free” synthetic community (SC) containing 61 isolates corresponding to 36 amplicon sequence variations (ASVs, identical 16S V5-V7 amplicon sequences) from the *Lj*SPHERE culture collection assembled from *Lotus* plants grown in Cologne soil [12]. This SC contains members from eight bacterial orders and 23 genera that represent a broad taxonomic diversity (Supplementary Table 6). This was used as a basic community to assess the impact of two critical properties of the symbiont: the capacity to produce Nod factor signaling molecules and to fix nitrogen on *Lotus* microbiota. For this, we assembled three additional communities by supplementing the SC with the wild-type symbiont *M. loti* R7A (SC+R7A), the Nod factor-impaired mutant *M. loti* R7A*nodC* (SC+R7A*nodC*), or the nitrogen fixation-impaired mutant R7A*nifH* (SC+R7A*nifH*). Wild-type *Lotus* plants were grown in a gnotobiotic system in the presence of one of these four bacterial consortia in two experimental conditions: with (3 mM KNO_3_ - see methods) or without inorganic nitrogen (Fig. 6a). The associated root and rhizosphere communities were analyzed after nine weeks by 16S rRNA amplicon sequencing (Fig. 6, Supplementary Fig. 11, and Supplementary Fig. 12). In the absence of inorganic nitrogen, only plants inoculated with R7A were nitrogen-replete because of nitrogen-fixing symbiosis. When nitrogen was present in the growth system, all plants developed shoots of similar weight regardless of the inoculum, indicating a nitrogen-replete condition (Supplementary Fig. 11a). Beta diversity analysis of bacterial communities using PCoA with Bray-Curtis dissimilarities identified a significant separation of root, rhizosphere, and input samples (Fig. 6b). Root microbiomes of plants exposed to communities containing R7A or R7A-*nifH* clustered together, irrespective of the experimental conditions. A second cluster contained root communities of plants inoculated with R7A*nodC* and SC as well as all rhizosphere communities, while the third cluster contained the inoculum communities (Fig. 6b). This separation indicates that root communities are primarily shaped by the presence of the plant and the presence of a Nod factor-producing symbiont, while rhizosphere communities are primarily shaped by the plant. A separate analysis only of rhizosphere samples identified that communities of plants inoculated with R7A separated from those exposed to other communities in both experimental conditions (Fig. 6c and 6d). This separation was driven by the symbiont and a reduced number of ASVs belonging to *Burkholderiaceae*, *Oxalobacteraceae*, and *Beijerinckiaceae* (ASV2, ASV22, and ASV30 in the nitrate-supplemented condition, and ASV32 in the absence of nitrate). In the absence of nitrate, we found that rhizosphere communities of plants inoculated with R7A*nifH* also separated from those with SC and R7A*nodC* (Fig. 6c). These results indicate that both Nod factor production and nitrogen fixation capacities of R7A impact the assembly of *Lotus* rhizosphere communities in nitrogen-deplete conditions. This was further confirmed in an independent experiment using a different, but proficient symbiotic *Mesorhizobium* strain isolated from Cologne soil (Supplementary Fig. 13) A further investigation at the taxonomic level identified that abundances of isolates belonging to seven families from Rhizobiales, Burkholderiales, Pseudomonadales and Xanthomonadales were significantly affected by the presence of a Nod factor-producing symbiont (Supplementary Fig. 11d, Supplementary Fig. 12a).

**Figure 6.**
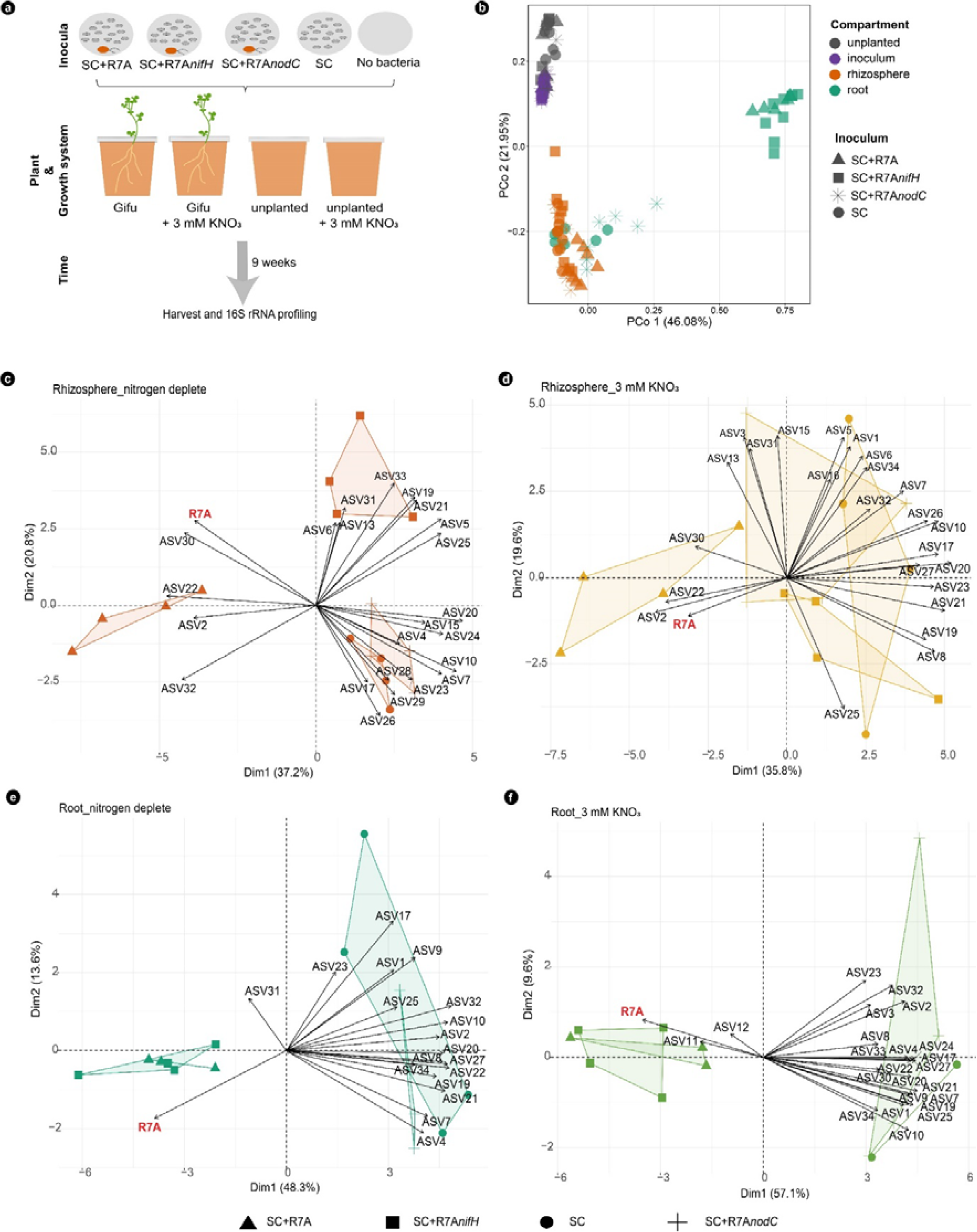
Nod factor-producing symbiont structures *Lotus* root-associated microbiota composition. **(a)** Experiment design. (**b**) PCoA analysis based on Bray-Curtis distances on all samples. Principal component analysis biplot of ASVs on rhizosphere (**c** and **d**) and root samples (**e** and **f**) from plants grown in the absence (**c** and **e**) or presence of KNO_3_ (**d** and **f**). ASVs in each sample are shown as variables. The arrows point out which condition is driven by the variable. The PCA plots are shown by the first and second dimensions.

Root communities of plants grown in these gnotobiotic conditions and in the presence of the Nod factor-producing *M. loti* were dominated by the symbiont (average 91%), irrespective of the presence or absence of nitrate or bacterial *NifH* gene (Fig. 6e, 6f and Supplementary Fig. 11e). This clear dominance of the symbiont is in marked contrast to the composition of root communities from plants grown with R7A*nodC*. Here, the relative abundance of the symbiont was reduced to 7.9%, and the root communities were enriched in members of Xanthomonadales, Burkholderiales, and Rhizobiales (Supplementary Fig. 11e, Supplementary Fig. 12b).

Together, these results obtained from studies based on gnotobiotic conditions and using tailored synthetic communities show that if symbiosis is enabled, the assembly of root and rhizosphere microbiota of *Lotus* is shaped by the presence of a Nod factor-producing symbiont, that provides the symbiont with competitive advantage for root colonization and affects the assembly of remaining members of microbiota, together providing a benefit for the plant host (Supplementary Fig. 13b).

### Exudates of *Lotus* plants exposed to the three nitrogen-dependent states vary in chemical composition

Root and rhizosphere microbiota are modulated by the exudates that plants secrete [14–16]. Thus, we hypothesize that *Lotus* plants grown in different nitrogen nutrition regimes will exude different metabolites. To test this, we grew *Lotus* plants in axenic, nitrogen starved or replete (10 mM KNO_3_) conditions, or in the presence of *M. loti* symbiont (Supplementary Fig. 14, material and methods). An additional condition where both the symbiont and the inorganic nitrogen were present was included in the assay to better differentiate between features produced in the two independent states. We detected a total of 1145 metabolite features (Supplementary Table 7) in the exudates collected from plants grown across the four conditions. As predicted, the composition and intensities of metabolites varied with the nitrogen-nutrition state (Fig. 7a), mirroring the separation observed in the root and rhizosphere microbiomes (Fig. 2b, c). We identified a similar number of metabolites across conditions, except for the starved plants where fewer features were detected (Fig. 7b). The metabolite features detected from these plants were generally of lower intensities compared to those detected in exudates from nitrogen-replete conditions (Fig. 7b and 7c), indicating that in the absence of nitrogen, *Lotus* plants maintained a diverse metabolite profile, but regulated their intensities. Large differences were detected for specific pathways of metabolites which varied across conditions in diversity and intensity (Fig. 7c). Carbohydrates and terpenoids, had significantly higher intensities in symbiotic plants (Fig. 7c), which is intriguing, since carbohydrates are recognized as chemical signaling and energy providers supporting microbial activities, while terpenoids are chemical compounds generally reported to be induced during plant defense and signaling [51, 52]. Next, we explored which metabolites were significantly enriched in the exudates collected from the three nitrogen dependent states (Fig. 7d). We identified 56, 384, and 416 features enriched in the exudates of plants grown in starved, symbiotic, or inorganic nitrogen conditions, respectively (Fig. 7d). Polyketides were overrepresented among features enriched in starved plants (5.6% compared to 0.25% in inorganic nitrogen and 0.6% in symbiotic states), while shikimates and phenylpropanoids were enriched in both starved and inorganic nitrogen conditions (13% in starved, 14% in inorganic versus 1.2% in symbiotic). Amino acids and peptides (31%), fatty acids (22%) and carbohydrates (8.8%) were more abundant in the symbiotic condition compared to the other two (Supplementary Table 7). An in-depth analysis of the 1145 features revealed that specific metabolites are significantly enriched in one of these conditions (Fig. 7e, Supplementary Fig. 15). Flavones, well-known attractants, and inducers of symbiosis signaling in rhizobia, as well as xanthones, also a phenolic metabolite, were enriched in starved plants. Jasmonic acids known as stress-related plant hormones were also enriched in the starved plants. Several metabolite classes with reported anti-microbial activities such as depsidone, amino acid glycosides, and oxygenated hydrocarbons were enriched in the exudates of plants grown in the presence of inorganic nitrogen. Metabolite classes enriched in the exudates of symbiotic plants were most diverse and varied from signaling and energy providing chemicals such as polysaccharides, disaccharides, fatty acyl glycosides to antimicrobial chemicals such as thiodiketopiperazine alkaloids and iridoids monoterpenoids.

**Figure 7.**
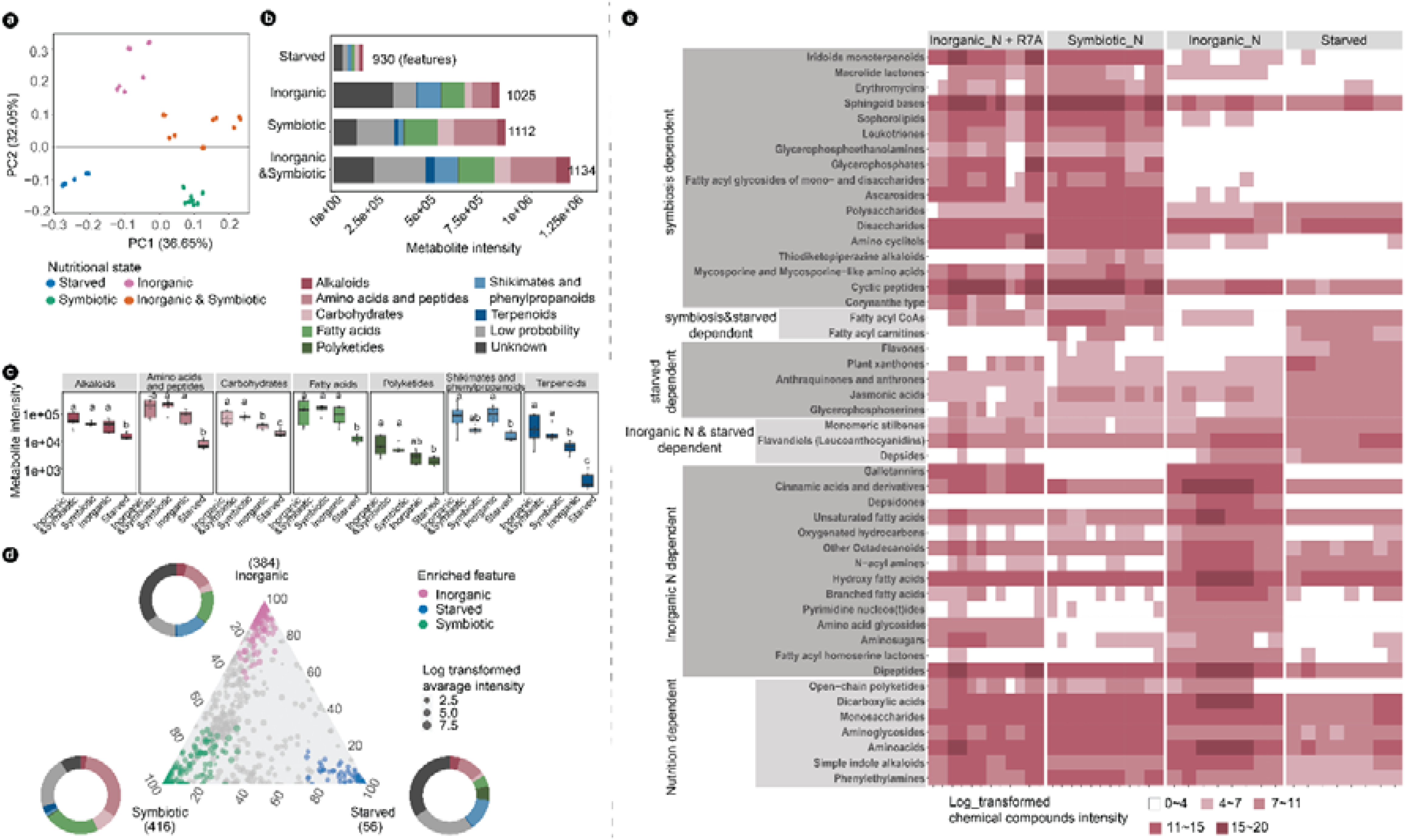
Distinct metabolites are identified in *Lotus* root exudates grown in symbiotic, starved, or inorganic nitrogen status. (**a**) PCA analysis of the chemical features identified in root exudates; Nitrogen nutritional status of samples are marked by colors; (**b**) Barplot illustrates the absolute abundance of chemical features in each of the nutritional status. The mean intensity of chemical features within samples is shown.; (**c**) Boxplot shows the abundance of metabolite intensity within the individual pathway, and statistical analysis is conducted within the individual pathway (*p*< 0.05); **(d)** Ternary plot illustrates statistically enriched features in the three nitrogen nutritional status, donut plots at each corner of the triangle shows the composition of metabolite intensity at the pathway level; **(e)** Heatmap shows enriched chemical compounds identified according to the nitrogen nutritional status. The chemical compound intensity in each of the sample is illustrated by color intensity.

Together, our analyses of metabolites revealed that *Lotus* plants adjust the panel and intensities of their root exudates according to a specific nitrogen regime. Importantly, we determined that plants grown in two different versions of nitrogen-replete conditions differ significantly in their exudate profile implying selection effects on the assembled bacterial communities.

## Discussion

Complex metabolic and signaling events are at the core of microbiota establishment in different environmental niches, and studies across environments and conditions suggest that a high level of taxonomic diversity is beneficial for microbiota homeostasis [53, 54]. Studies in human, mice, zebrafish, and fly have shown that perturbations in host diet have direct and predictable consequences on their gut microbiota [55–59]. The soil provides nutrients required for plant growth and its properties have the largest impact on rhizosphere and root microbiota [11, 60, 61]. In addition to soil, plant pathogens can directly modulate the microbiota through secretion of antimicrobial effectors [62]. Here, we investigated the role of symbiont-induced signaling on the assembly of *Lotus japonicus* root microbiota. Symbiosis between nitrogen-fixing bacteria and legumes was found to contribute to a diverse and beneficial root-associated microbiota [7, 42, 43, 45]. The current study provides evidence that symbiont-derived Nod factor signals contribute to microbiota homeostasis by inducing the Nod factor-dependent signaling in the host, which in turn ensures a diverse and interconnected bacterial microbiome (Fig. 8). Results from analyses using bacterial mutants as well as different plant genotypes grown in nitrogen-deplete or replete conditions revealed a tight dependency of *Lotus* microbiota on the capacity of the host to mount an active Nod factor signaling. Root microbiota of soil-grown plants was gradually affected, according to the degree of impairment in the Nod factor signaling present in the three mutants (Fig. 1b). A possible explanation for this pattern is that Nod factor signaling leads to changes in the physiology of the plant root, which in turn impact the association with commensal members. Indeed, we found that the chemical profile of exudates from symbiotically active plants differs significantly from that of starved or inorganic nitrogen-supplemented plants (Fig. 7a, 7d and 7e). The symbionts present in the soil were not identified as hubs by our network analyses but emerged as major players in structuring the communities indirectly, via a host-mediated feedback effect on the remaining members of bacterial communities. A similar feedback control of microbiota was previously described for anaerobic, beneficial commensals producing short-chain fatty acids (SCFA) during colonization of mice colon [55]. SCFAs activate PPAR-γ-signaling in mice, which ensures a metabolism that preserves hypoxia in the colon environment, and thus maintains microbial homeostasis limiting the expansion of members from *Enterobacteriacea* that induce dysbiosis [63]. The *nfr5* and *chit5* plants with impaired Nod factor signaling, as well as wild-type plants inoculated with SC+R7A*nodC* or SC, are starved and have associated communities that differ from those of symbiotic plants (Fig. 2, Fig. 3, and Fig. 6). Interestingly, we found that in controlled reconstitution experiments with an absent Nod factor signaling, there is an increased abundance of members from Pseudomonadales and Xanthomonadales (Supplementary Fig. 11e). The isolates included in our studies are commensals, however well-described plant pathogens emerge from these two taxa [64, 65], thus an increased abundance could provide increased chances for pathogenic associations to evolve or become apparent. Future studies may reveal whether Nod factor signaling limits root dysbiosis in legumes.

**Figure 8.**
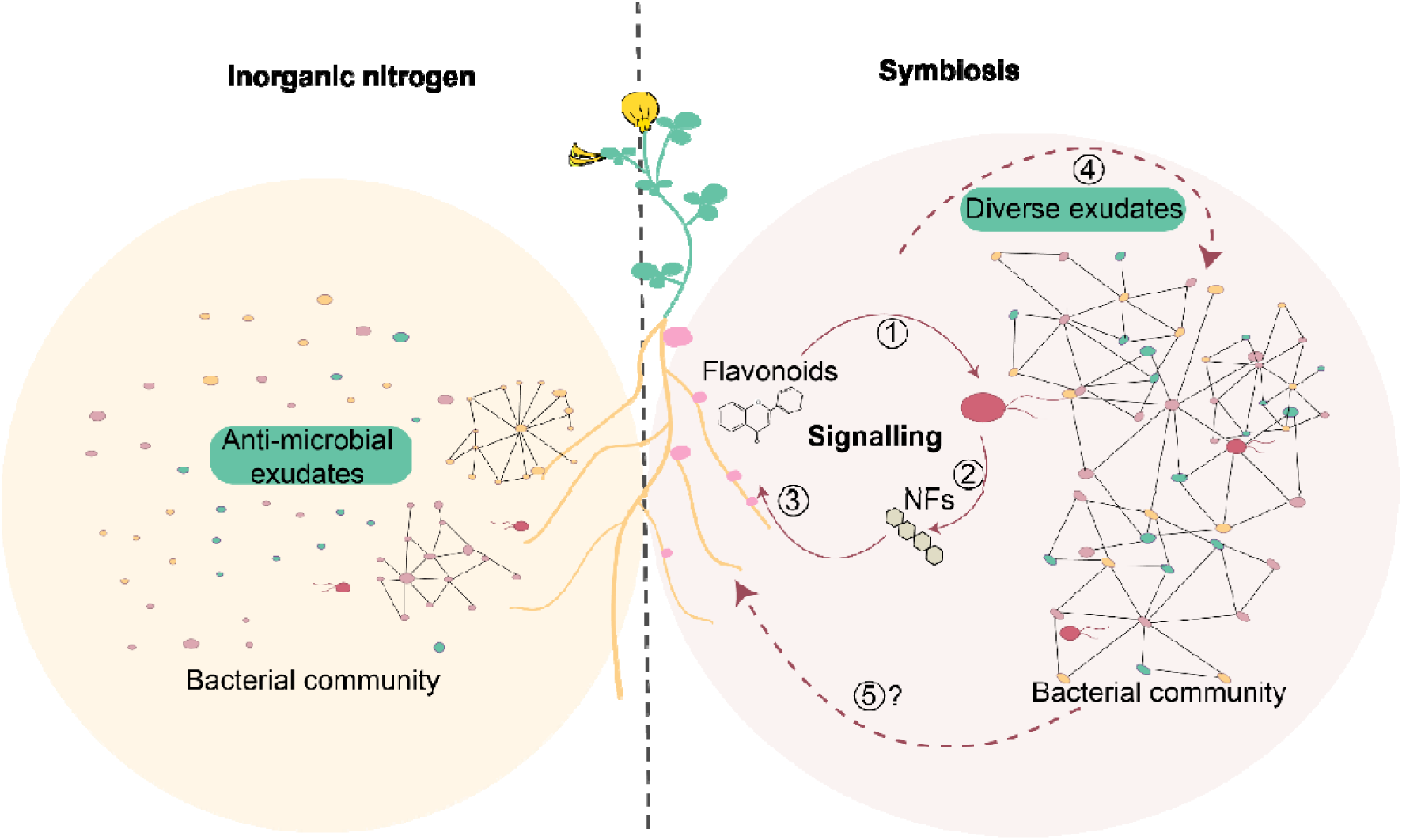
Nitrogen nutrition and signaling during root nodule symbiosis impact the community assemblies. *Lotus* plants grown in the presence of inorganic nitrogen secrete specific metabolites and assemble a microbial community withlow connectivity. *Lotus* plants grown in symbiosis permissive conditions secrete metabolites such as flavonoids (1) that induce Nod factor production in compatible nitrogen-fixing Rhizobium isolates (2). Nod factors are recognized by the *Lotus* host which initiates a signaling pathway (3) to accommodate the symbiont. Symbiotically active roots have an exudate profile (4) and associated microbial communities that differ from plants grown in the presence of inorganic nitrogen. It remains to be determined how bacterial communities associated with symbiotically active plants impact the host to promote the symbiotic association and plant growth (5).

The role of *Lotus* host on structuring the microbiota is crucial, knowing that Nod factor production is induced in symbionts primarily when host-specific (iso)flavonoids are secreted from starved roots [66]. Furthermore, the host controls the activation of Nod factor signaling in the root only when compatible Nod factors are perceived by specific LysM receptors [67]. Thus, the *Lotus* host is in continuous control of its colonization by the symbiotic partner and of its associated microbiota. This is in line with recent results emerged from evolutionary modelling studies predicting that cooperation in host microbiome is driven by the host control over symbionts [68, 69].

We show here that microbiota associated with *Lotus* plants provided with different sources of nitrogen (inorganic or symbiotic) are significantly different. *Lotus* plants found in a starved, symbiotic, or inorganic nitrogen-replete state differ in their root and shoot transcriptomes [70], metabolic state[71] and exude different compounds in the rhizosphere (Fig. 7). Here we find that these states are associated with distinct microbiomes and that particular genera enriched in each of the states have a high level of predictability (Supplementary Table 5). The inorganic nitrogen induced significant changes in the soil, rhizosphere, and root communities of *Lotus*. A similar response was reported for wheat and tomato [72, 73]. Importantly, analyses of the co-occurrence networks for microbiomes present in the three states of *Lotus* plants revealed the clear negative effect of the inorganic nitrogen on microbial connectivity. At this stage, it is unclear whether this is a result of changes in the host metabolism and root exudate pattern (Fig. 7), a result of the functional capacities exerted by the members of these communities, or both.

Collectively, our findings provide evidence that signaling established between the plant host and the compatible symbiont impacts the assembly and properties of the remaining members of the community. Furthermore, it supports the idea that Nod factor signaling established in legumes after recognition of nitrogen-fixing bacteria facilitates not only nitrogen nutrition, but also the association with diverse and highly connected bacterial communities at the root-soil interface supporting nitrogen-fixing symbiosis.

## Materials and Methods

### The plant material and growth in soil

The ecotype Gifu B-129 of *L. japonicus*, and the symbiosis-deficient mutants *nfre:1*, *nfre:2* [40], *chit5:1*, *chit5:2* [41], *nfr5:2*, and *nfr5:3* [29] were grown in Cologne Agriculture soil (CAS9) under greenhouse conditions (day/night cycle = 14/10 h, temperature: day/night = 22/18°C, humidity: 75% Rh). Plants were watered either by sterile water or 10 mM KNO_3_, and harvested at 9 weeks old.

### Plant growth in gnotobiotic system for microbiota reconstitution

Magenta boxes filled with clean LECA were used as the gnotobiotic system. Individual liquid cultures of bacterial isolates (Supplementary Table 6) were washed twice by centrifuging at 4700 rpm for 15 minutes, discarding supernatant and resuspending with sterile water. The washed bacterial liquid cultures were pooled together accordingly. OD_600_ of the SynComs was adjusted to 0.02. The magenta boxes were applied by 50 ml SynComs (OD_600_=0.02), and 10 plants were potted in it, sitting in 21°C with a 16/8 light/dark cycle condition. The plants were kept in closed magenta boxes for the first four weeks, and then the lids of magentas were removed and the ¼ B&D media (with or without 3 mM KNO_3_) was supplied to the plants regularly. Note that our previous experiment using 10 mM KNO_3_ in this gnotobiotic system had a negative impact on plant growth, thus the concentration of KNO_3_ were reduced to 3mM. Plants were harvested at 9 weeks old.

### Plant growth in gnotobiotic system for collecting root exudate

Gifu wild-type plants were grown on sterile Petri dishes filled with ¼ Broughton and Dilworth agar media supplemented or not with 10 mM KNO_3._ Unplanted plates were included as control. The symbiotic state was reconstituted by inoculation of Gifu plants with *M. loti*, R7A. Root exudates were collected after 4 weeks of growth.

### Sample collection and 16S rRNA amplicon sequencing

At the harvesting stage, fractionation was conducted on the root system (4 cm-long starting 1 cm below hypocotyl), and rhizosphere, root, and nodules were separated by wash process (three times sterile water, two times detergent) and surface sterilization (one time 80% ethanol, one time 3% bleach). Nodules and visible primordia were separated from root fragments under microscope using a scalpel. The collected root, rhizosphere, nodules, as well as soil/LECA samples were homogenized, and DNA was extracted using the FastDNA Spin kit for Soil (MP Bioproducts) according to the manufacturer’s protocol. The variable V5–V7 region of 16S were amplified using primer pairs 799F (AACMGGATTAGATACCCKG) and 1192R (ACGTCATCCCCACCTTCC) at the first round PCR. Barcode primers for distinguishing samples were targeted at the 1192R and sequencing adapters of Miseq at the second round PCR. The indexed 16S rRNA amplicons were pooled, purified and sequenced by Illumina Miseq platform.

### Preprocessing of raw reads for 16S rRNA amplicons

Raw reads of 16S rRNA amplicons were processed using a method that combined QIIME [74] and USEARCH [75]. ASV clustering was conducted using the UNOISE algorithm [76]. Reads that had 97% identity to clustered ASVs were mapped as read counts to generate the ASV table. Taxonomy assignment was done with SINTAX algorithm. The AsCoM reference database developed on CAS soil was used as reference for taxonomy assignment [49]. ASVs assigned as mitochondrial or chloroplast and which had relative abundance less than 0.01% across all the samples were removed. The filtered ASV table was used for downstream analysis.

The raw reads obtained from the reconstitution experiment were preprocessed by mapping reads back to reference sequences (USEARCH, UPARSE-REF algorithm). Reads that matched 99% or above to reference sequences were kept for generating the ASV table and for downstream analysis.

### Sample collection and analysis of root exudate

Chemicals exuded by the roots or present in the control unplanted plates onto sand grains after O/N exposure were collected by sterile water wash (Supplementary Fig. 14) and went through syringe filters with 0.45 µM pore size. The collected samples were analyzed by an ultra-high-performance liquid chromatograph (UHPLC) coupled to a quadrupole time-of-flight mass spectrometer (qToF MS, Bruker Compact) with electrospray ionization. Untargeted analysis was performed. The raw data were preprocessed by MzMine 3.3.0 [77] (Supplementary Fig. 14) to construct the feature list. Features detected at the end of the chromatogram (retention time > 19 mins) were discarded. Chemical classes were assigned to metabolites with the NPC compound classification system by SIRIUS 5.6.3 [78]. Detailed parameters used for the feature list construction can be found on uploaded MzMine batch file (MassIVE, user: MSV000092000, password: lotusdataset123).

### Statistical analyses and data visualization

The statistical analyses and most data visualization were conducted in R v4.2.3.

### Alpha and Beta diversity

The ASV table was rarefied according to the lowest read numbers in the samples to calculate the Chao1 diversity indices. Significant differences were determined using the Kruskal-Wallis test (krus.test in R, p<0.05). To estimate beta diversity for the samples, the ASV table was normalized using the cumulative sum scaling method [79]. Bray-Curtis distances between samples were used for constrained or non-constrained principal coordinate analysis (CPCoA, capscale function; PCoA, cmdscale function). The PERMANOVA analysis was performed by the adonis2 function from the vegan package in R.

### Differential abundance

The R-package edgeR v3.36.0 was used to fit a negative binomial regression with quasi-likelihood estimation [80] to the ASV counts. The null-hypothesis |log2FC| ≥ 1 was tested with the glmTreat function [81]. Correction for multiple testing was carried out using the Benjamini-Yekutieli adjustment [81]. The nominal FDR-level was set to 0.05. The ternary plots were constructed using ggtern v3.3.5 [82].

### Phylogenetic tree, heatmap and visualization

The ASV sequences were aligned by clustalo [83] and used to construct phylogenetic trees by raxml [84]. The phylogenetic trees were adjusted according to published phylogeny [12]. The RA of ASV were log2 transformed for heatmap visualization using the Interactive Tree of Life web tool [85].

### Co-occurrence networks

Correlations between ASV abundances were estimated using fastspar [86]. The correlation strength exclusion threshold for SparCC’s iterative procedure was set to 0.2. Testing for significant correlations was performed using the permutation bootstrapping procedure implemented in fastspar (20,000 for each condition, 50,000 specifically for rhizosphere in the inorganic nitrate state). The resulting p-values were corrected for multiple testing using Benajmini-Hochberg with a nominal FDR-level of 0.1. To equalize the means between genotypes, the function “removebatcheffect” from the package limma v3.50 [87] was used. The correlations with absolute value that less than 0.6 were discarded. The correlation networks were visualized in Gephi, and the modularity, betweenness centrality, and degree of correlation were calculated based on the correlations between ASVs.

### Prediction analysis

Prediction of the nitrogen status was carried out on the data from Zgadzaj et. al. [7] using genera as predictors. Candidate covariates were identified with differential abundance analysis after collapsing the ASV data from the soil experiment into genera. The Zgadzaj et. al dataset was split into a training and test dataset. A third of the biological replicates from each nutritional status were assigned to test data and the rest was assigned to the training data. Independent filtering was performed by removing any genus that did not appear in at least 90% of samples. The centered log-ratio (CLR) transformation was applied to the genus abundances. Variable selection was carried out on the training data using the procedure described in Diaz-Uriarte et al. [50]. A support vector machine was fitted on the training data with the covariates selected during variable selection. The predictions were validated by applying the model to the test data.

### Identifying metabolites with differential abundance in the different nutritional states

The metabolite data was normalized using a procedure similar to the default option in DESeq2 [86]. After the normalization, zeros were replaced by 0.5 and the data was log-transformed. Metabolites present in less than 10% of the samples were not included in the analysis. Next, we applied a linear mixed model [89, 90] to identify metabolites where the effect of the nutritional state differ significantly between the Gifu samples and the control samples. Only the metabolites with p<0.05 after adjusting for multiple testing with Benjamini-Yekutieli were retained for further analysis. Then, pair-wise tests between the states were conducted on the Gifu samples using another linear mixed model, and p-values were corrected for multiple testing with Benajmini-Hochberg.

### Data and code available

All the scripts used for this study have been deposited in the github(https://github.com/taoke1/Project_of_nitrogen_source_and_Nod_factor_signalling). All the amplicon raw data has been deposited in the National Center for Biotechnology Information (NCBI) database with accession number PRJNA974421. The metabolomic raw data files are stored as a MassIVE dataset (user: MSV000092000, password: lotusdataset123).

## Acknowledgments

We thank Prof. Paul Schulze-Lefert and Dr. Ruben Garrido-Oter_ for their continuous support of our studies and critical reading of the manuscript. We thank Dr. Kathrin Wippel for constructive discussions and critical reading of the manuscript. We thank Dorthe B. Jensen for helping with the experimental set-up and Finn Pedersen for propagation of plants. We thank Taylor Grace FitzGerald for proofreading the manuscript.

## Funding

This work was supported by the Bill and Melinda Gates Foundation and the UK’s Foreign, Commonwealth and Development Office (FCDO) through the Engineering Nitrogen Symbiosis for Africa project (ENSA; OPP11772165), the Danish Council for Independent Research (9041-00236B), the Molecular Mechanisms and Dynamics of Plant-microbe interactions at the Root-Soil Interface project (InRoot), supported by the Novo Nordisk Foundation grant NNF19SA0059362. The China Scholarship Council supported K.T. and S.Z. for their Ph.D. study.

## Author contributions

K.T., S.Z., and A.M. performed experimental studies of the plants with help from Z.B. and S.K.. I.T. and K.T. performed computational analysis of the data. C.S. and P.T. performed chemical analyses of root exudates. E.V.R., I.T., and K.T. performed statistical analyses of root exudate metabolites. S.R. and K.T. conceived the experiments, S.R., M.G. and R.W. coordinated studies, K.T, I.T, and S.R. wrote the manuscript with input from all authors.

## Competing interests

The authors declare no competing interests.

## Notes

### Competing Interest Statement

The authors have declared no competing interest.

