## Supplemental Figures for "Nitrogen source and Nod factor signaling map out the assemblies of *Lotus japonicus* root bacterial communities"

**Supplementary Figures**

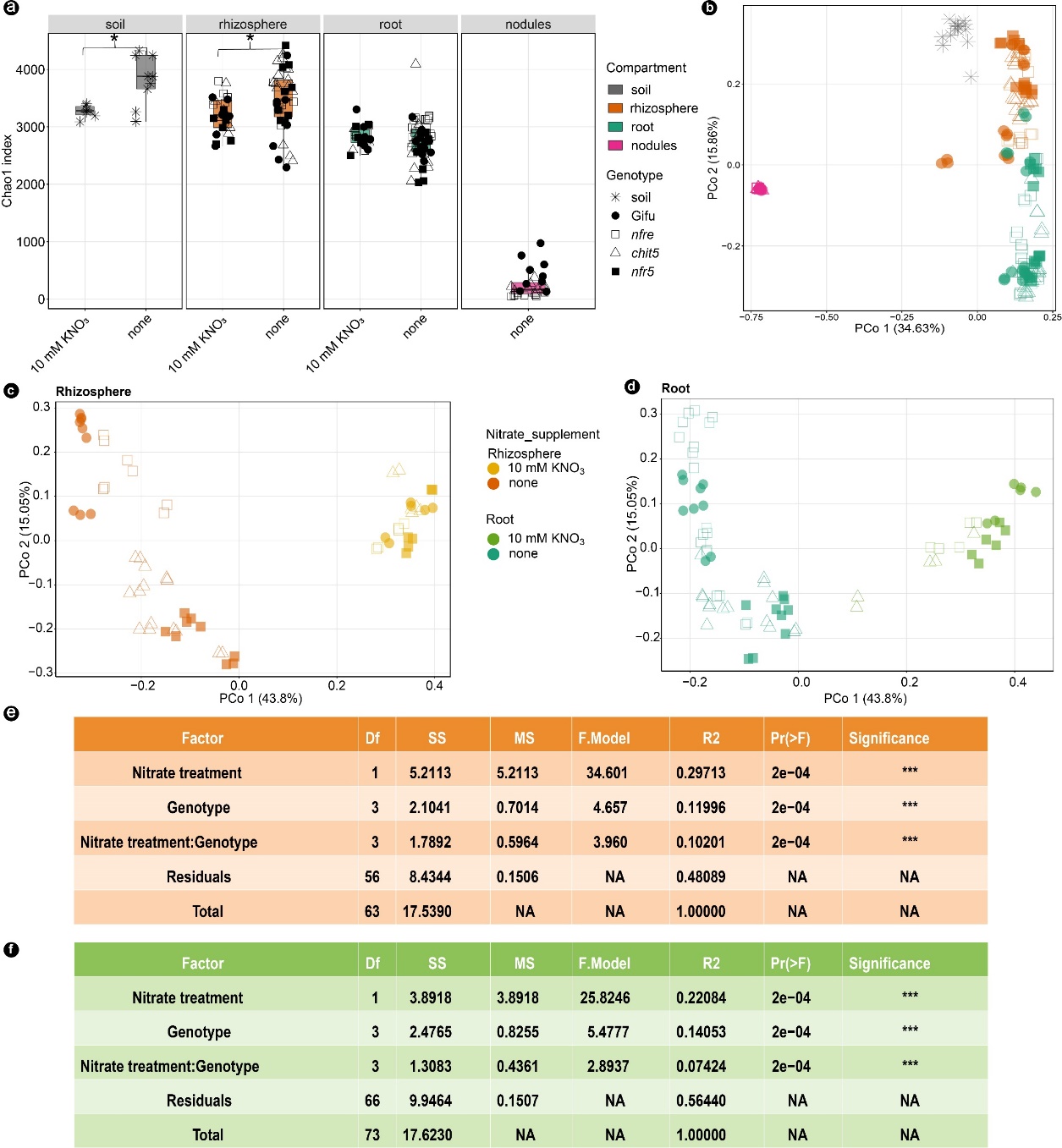

**Supplementary Figure 1 Diversity analysis for bacterial communities associated with roots of Gifu, *nfre*, *chit5*, and *nfr5* grown in Cologne soil.** a) Chao1 index for communities from soil, rhizosphere, root, and/or nodules compartments. Mann-Whitney U-testdetects significant differences between samples from native or nitrate-supplemented conditions. PCoA plot of Bray-Curtis distances including b) all the samples, rhizosphere c) and root d) samples. PERMANOVA on rhizosphere e) and root f) samples illustrate that nitrate treatment is the main driver for rhizosphere community, while this influence is reduced for the root samples where the effect of the genotype is larger.MS: mean of square; SS: sum of square; Significance Codes: 0 ‘***’ 0.001 ‘**’ 0.01 ‘*’ 0.05 ‘.’

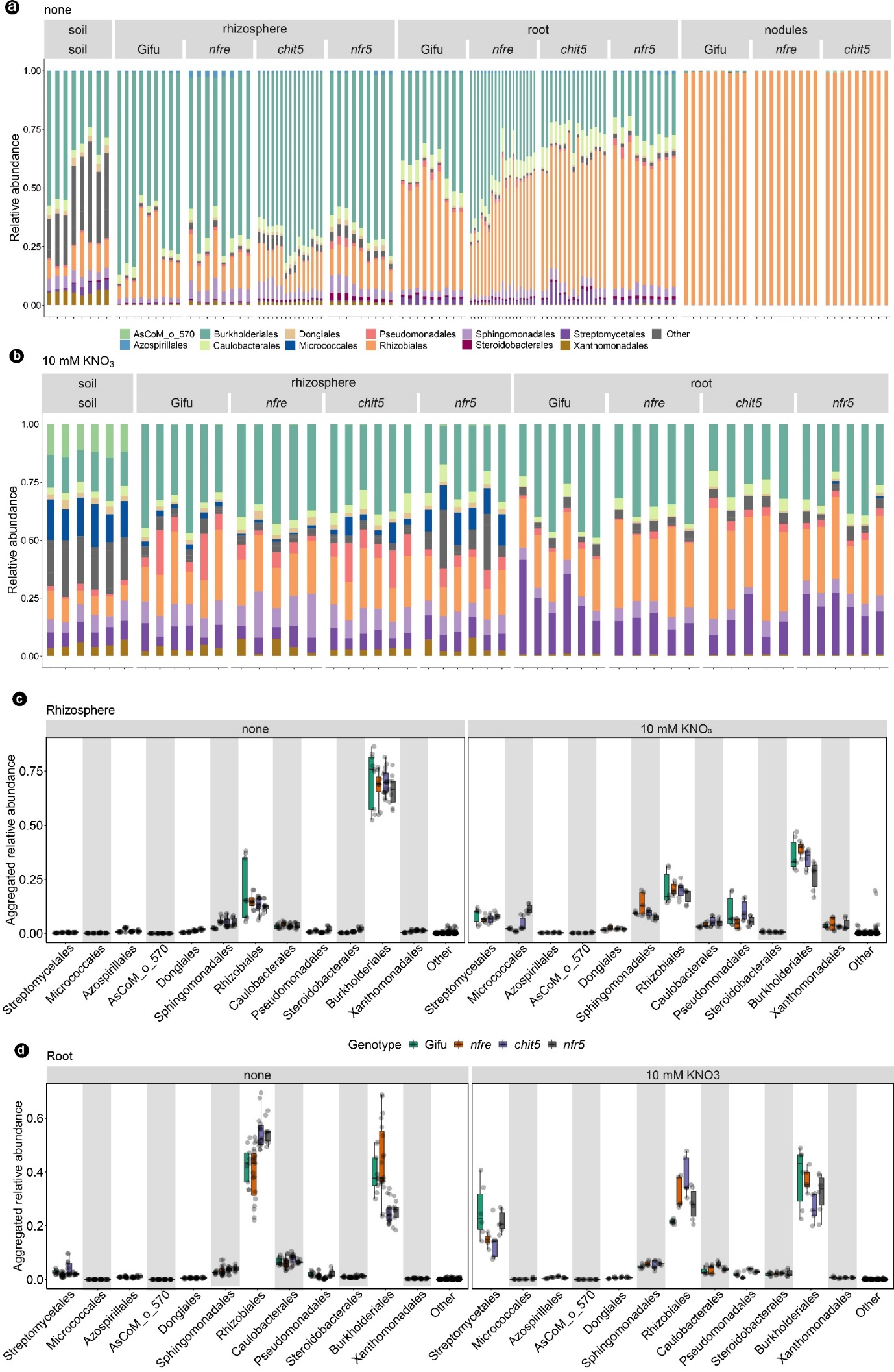

**Supplementary Figure 2. Relative abundance of bacterial orders in bulk soil, rhizosphere, root, and nodules samples collected from Cologne soil supplied with sterile water (a and c) or with 10 mM KNO_3_ (b and d).** In the stack bar plot (**a** and **b**), columns indicate the replica and colors indicate taxonomic assignment. In the boxplot (**c** and **d**), colors indicate genotypes.

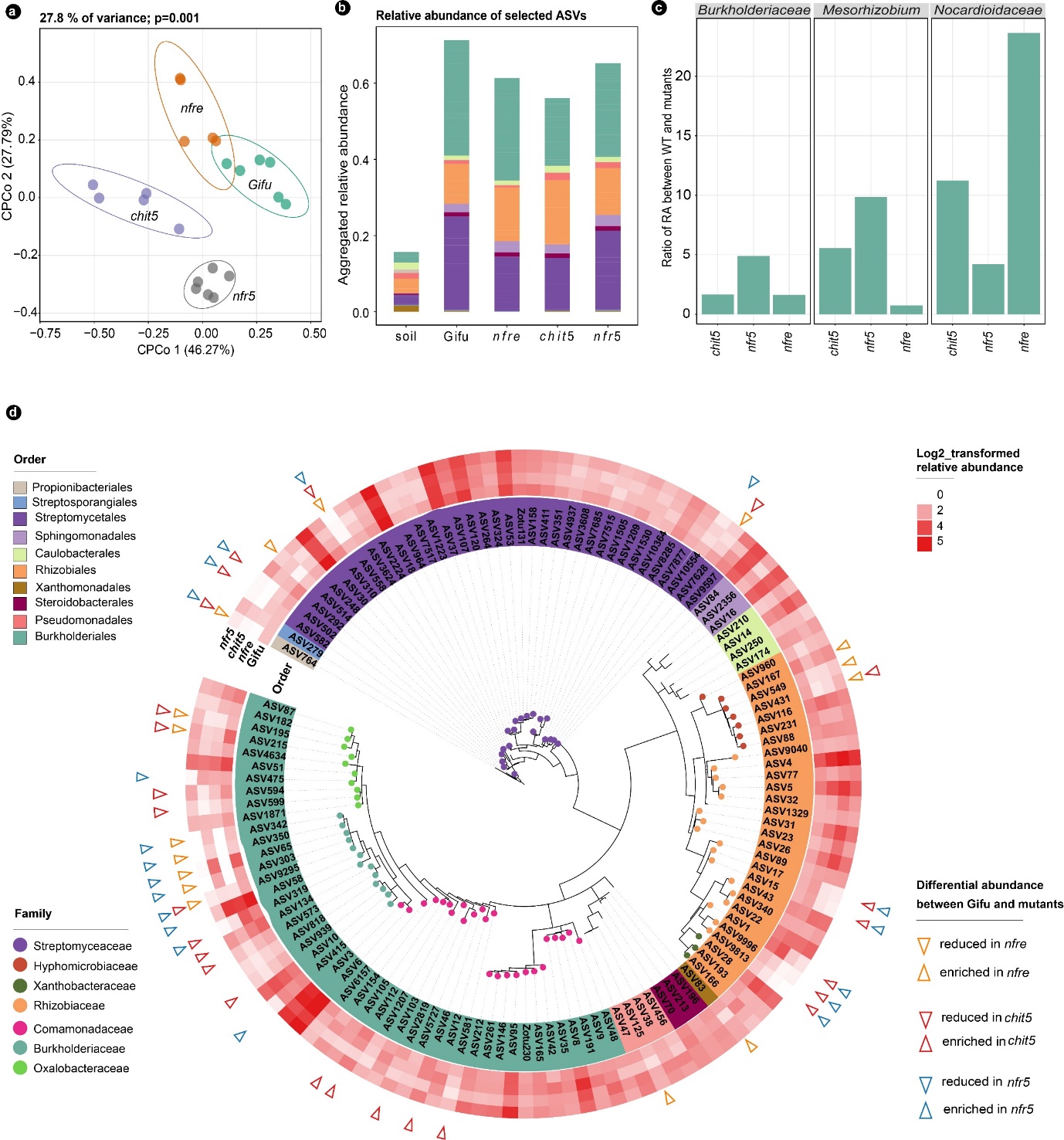

**Supplementary Figure 3. Impairment of Nod factor signaling changes the composition of root bacterial community of *Lotus* plants grown in nitrate-supplemented Cologne soil. a**) Communities associated with roots of wild-type, *nfr5*, *nfre*, and *chit5* are distinct and separated from each other. **b)** Cumulative relative abundance of selected ASVs (RA >0.3% in roots of Gifu) in soil and roots of the four genotypes. **c)** The ratio of RA between Gifu and mutants of the top three taxa in the roots based on selected ASVs: RA>0.3% in roots of Gifu. **d)** Distinct ASVs have a significantly different RA in mutant roots compared to wild-type Gifu. ASVs with RA > 0.3% in roots of Gifu are presented in a phylogenetic tree constructed based on the 16S rRNA V5-V7 region. The taxonomic information is shown by color on the name of the ASV (order) and by color on the tree branch (family). The heatmap shows the log2-transformed RA of each ASVs in the roots of Gifu, *nfre*, *chit5*, and *nfr5* plants. Empty triangles on the outer layer of the heatmap point out ASVs that potentially have a different abundance compared to wild-type plants.

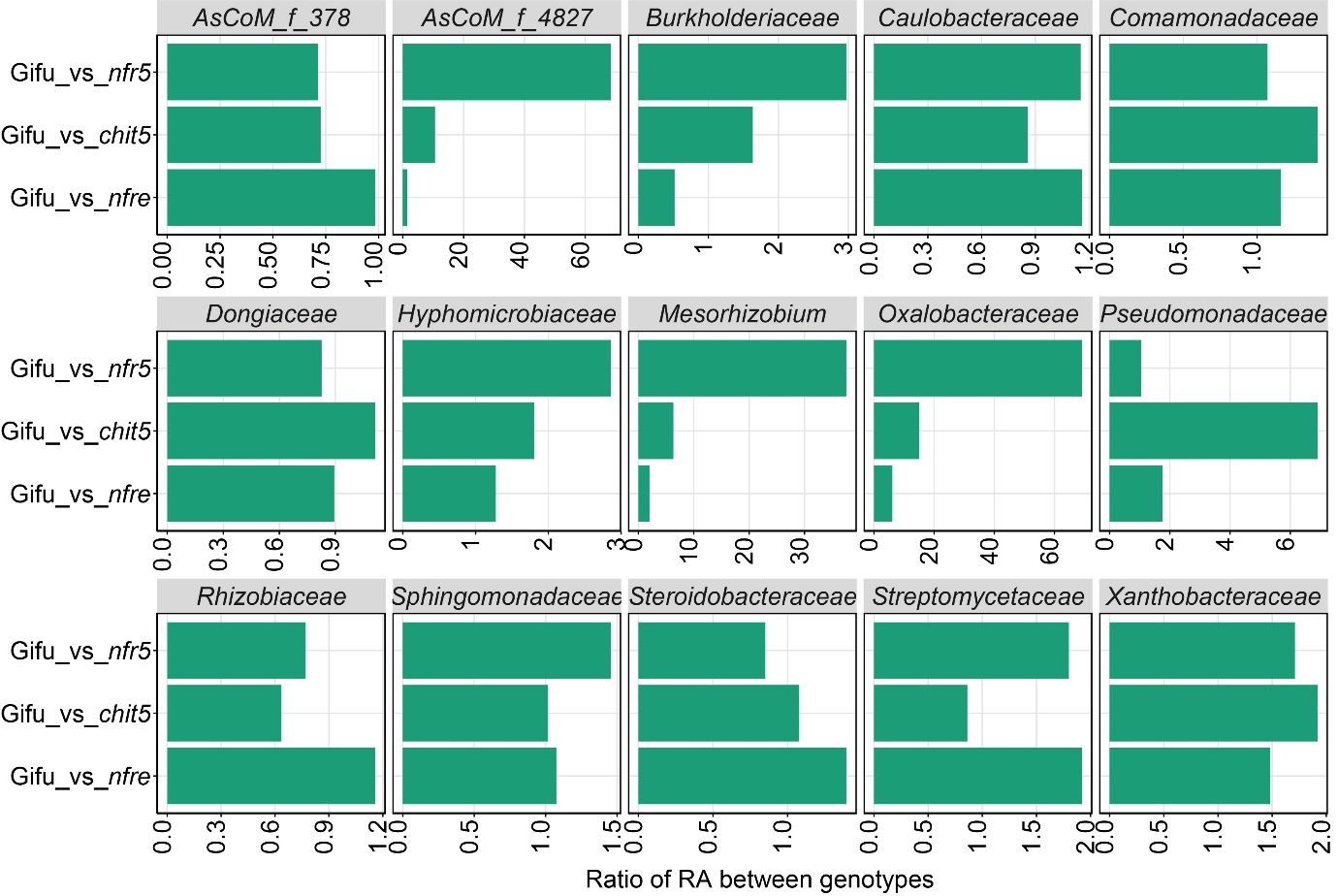

**Supplementary Figure 4. The ratio of RA for families between wild-type and mutants at the taxonomic family level on roots grown in water supplied condition.** The calculation of ratio is based on selected ASVs (RA>0.3% in roots of Gifu).

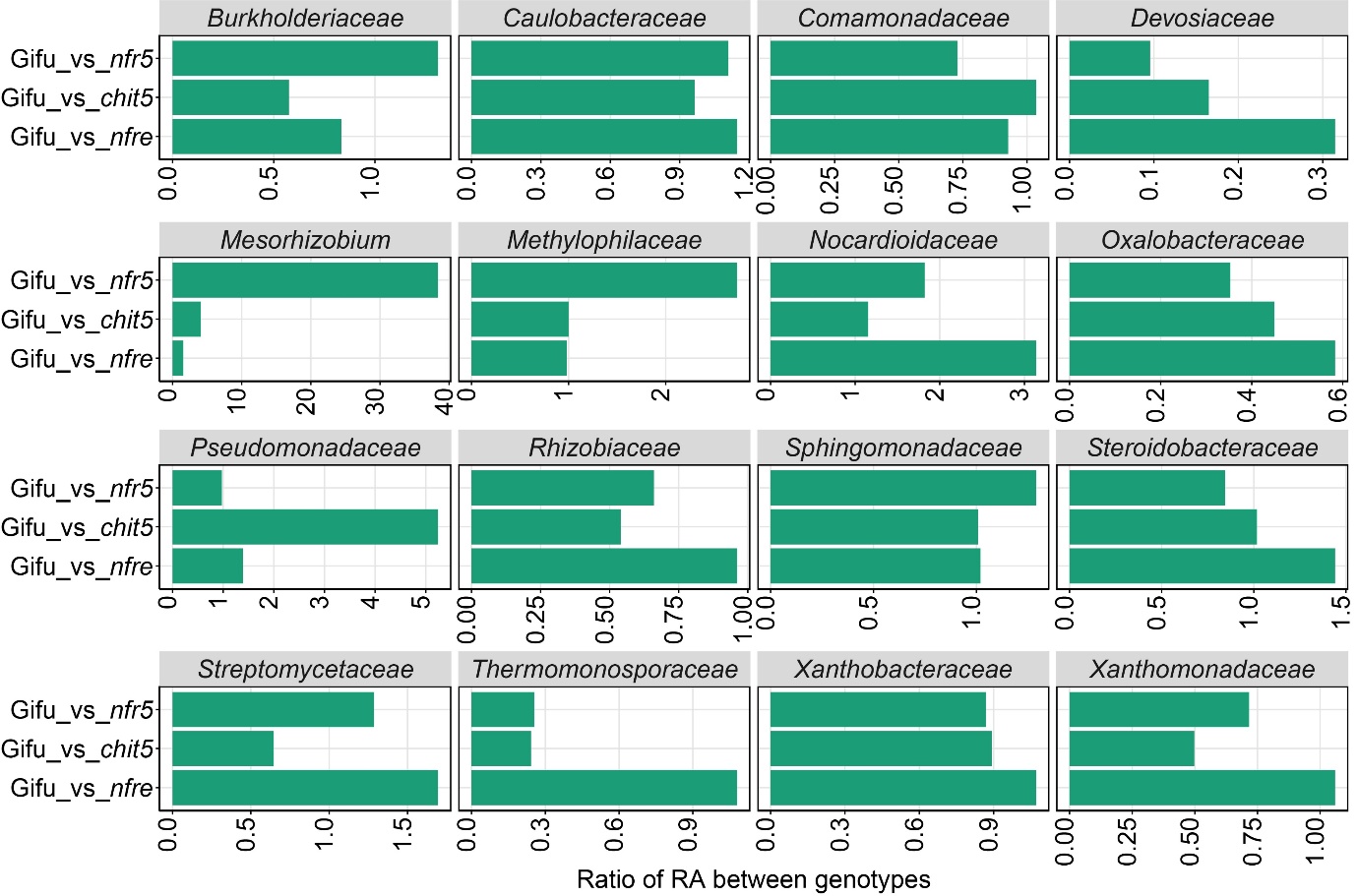

**Supplementary Figure 5. The ratio of RA for selected families between wild-type and mutants at the taxonomic family level on roots grown in 10 mM KNO_3_ supplied condition.** The calculation of ratio is based on selected ASVs (RA>0.3% in roots of Gifu).

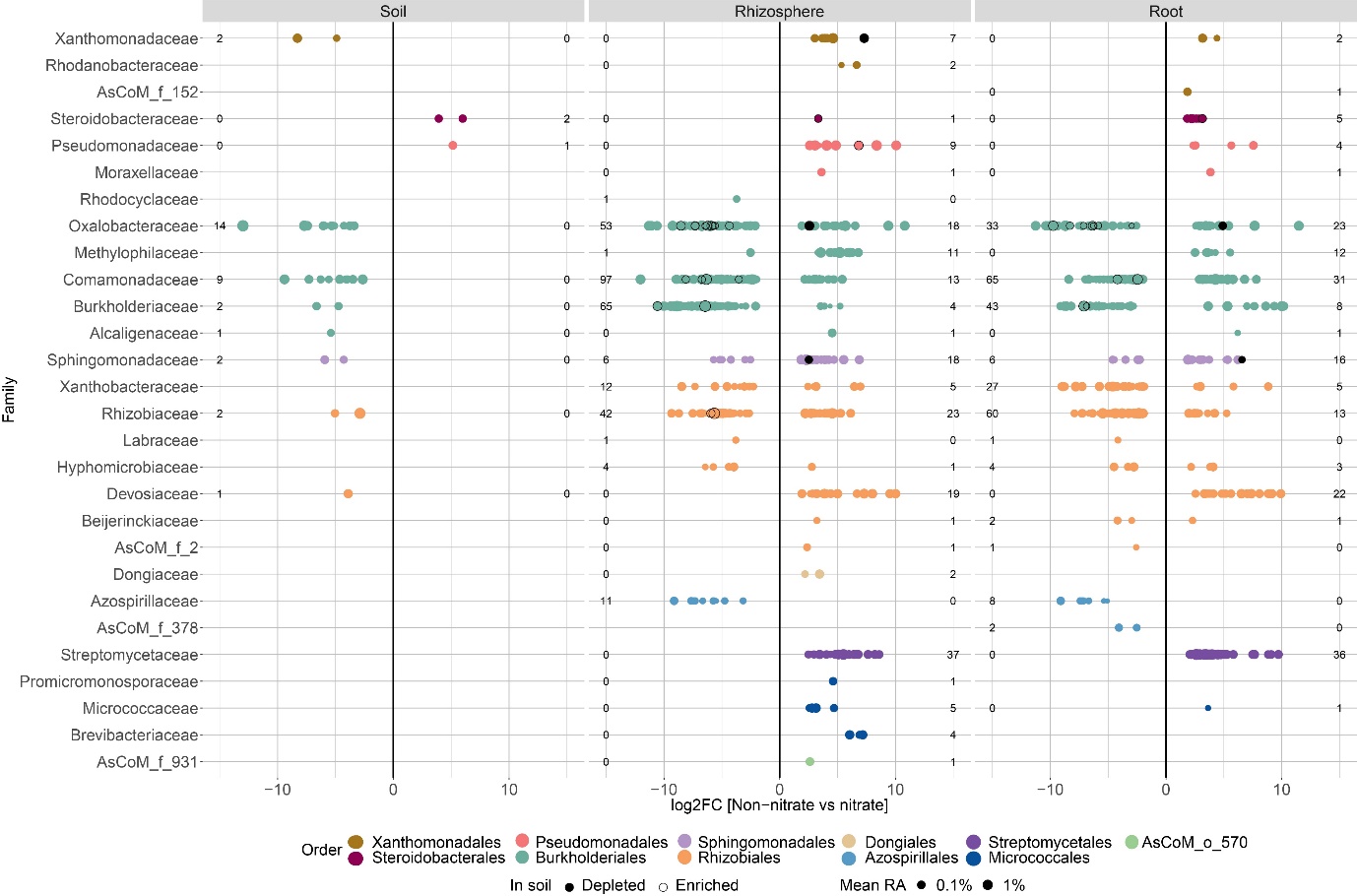

**Supplementary Figure 6. ASVs belonging to different bacterial taxa are differentially enriched in soil, rhizosphere, or roots of Gifu plants when grown in native or nitrate-supplemented Cologne soil.** Each dot presents an ASV significantly different in abundance in nitrate versus native soil conditions. The size of the dot represents the relative abundance in the condition in which it is enriched. The color of the dots corresponds to the taxonomic order assignment. The numbers at the left/right edge of the plots represent the number of ASVs within the respective family found to be significantly different between conditions. The black dots and dots with black outlines in rhizosphere and root samples indicate ASVs that were also differentially abundant in the nitrate versus native soil. These dots indicate ASVs having a different (black) or similar (black outline) pattern of enrichment/depletion as observed in the soil.

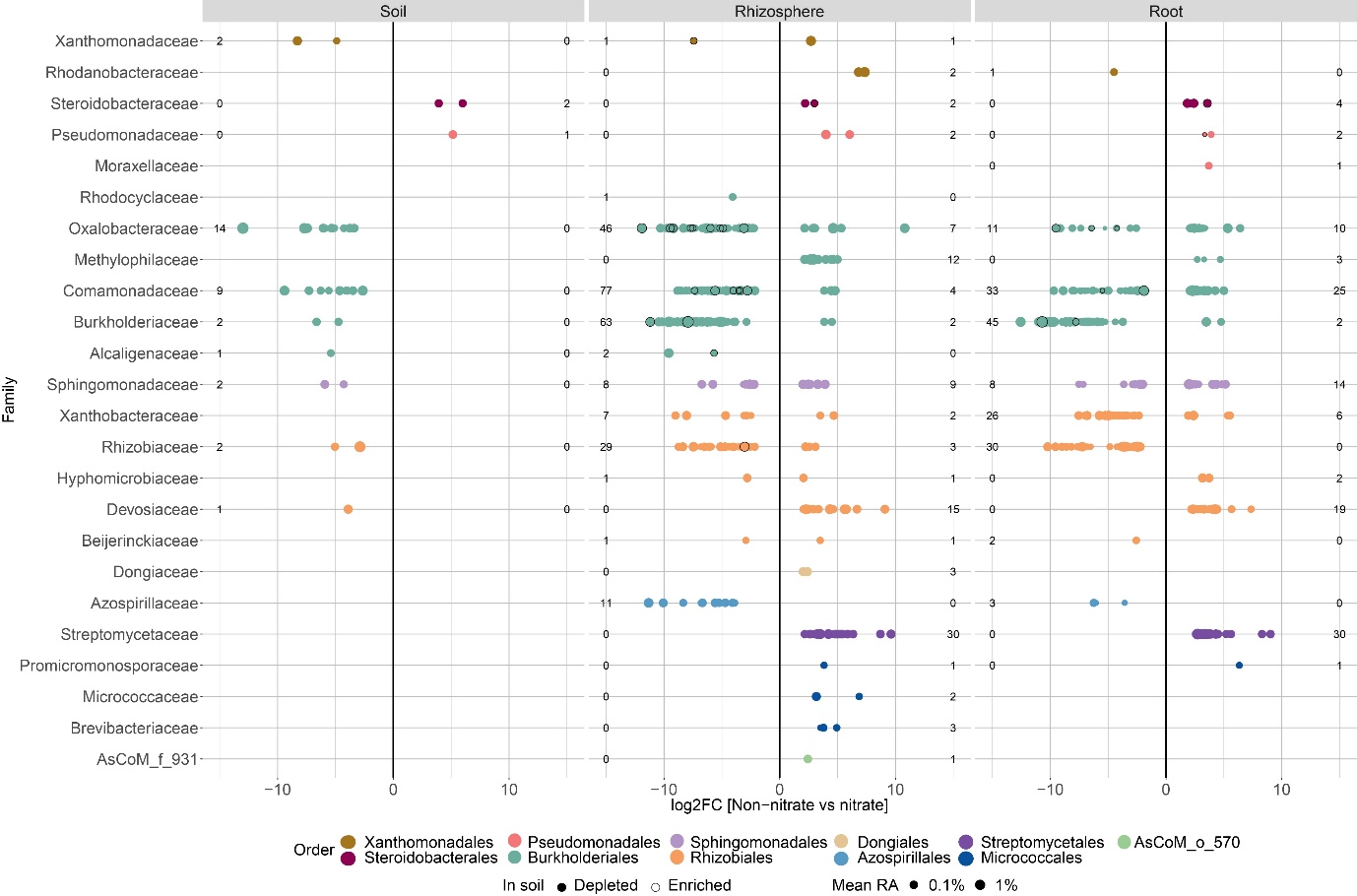

**Supplementary Figure 7. ASVs belonging to different bacterial taxa are differentially enriched in soil, rhizosphere, or roots of *nfre* plants when grown in native or nitrate-supplemented Cologne soil.** Each dot presents an ASV significantly different in abundance in nitrate versus native soil conditions. The size of the dot represents the relative abundance in the condition in which it is enriched. The color of the dots corresponds to the taxonomic order assignment. The numbers at the left/right edge of the plots represent the number of ASVs within the respective family found to be significantly different between conditions. The black dots and dots with black outlines in rhizosphere and root samples indicate ASVs that were also differentially abundant in the nitrate versus native soil. These dots indicate ASVs having a different (black) or similar (black outline) pattern of enrichment/depletion as observed in the soil.

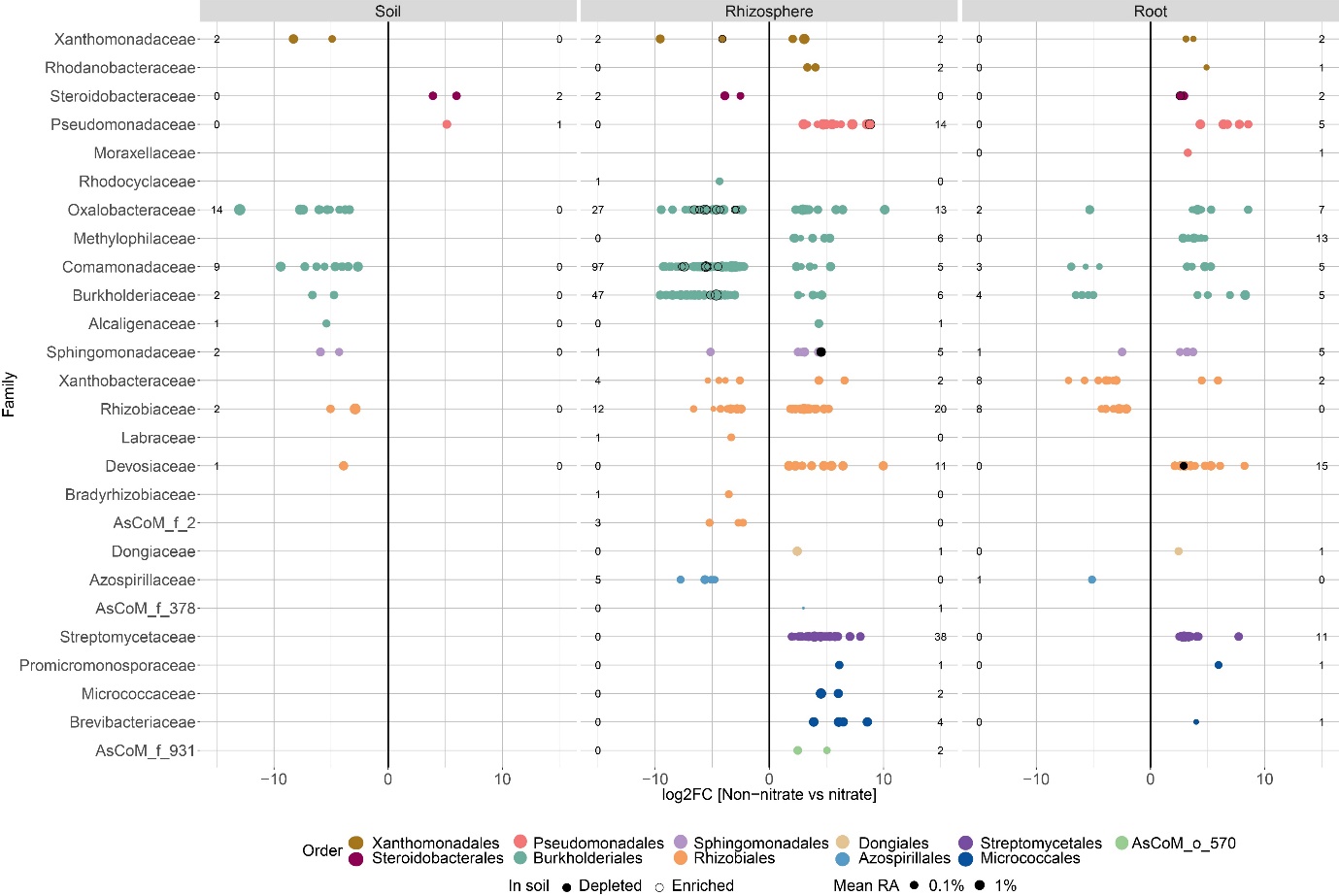

**Supplementary Figure 8. ASVs belonging to different bacterial taxa are differentially enriched in soil, rhizosphere, or roots of *chit5* plants when grown in native or nitrate-supplemented Cologne soil.** Each dot presents an ASV significantly different in abundance in nitrate versus native soil conditions. The size of the dot represents the relative abundance in the condition in which it is enriched. The color of the dots corresponds to the taxonomic order assignment. The numbers at the left/right edge of the plots represent the number of ASVs within the respective family found to be significantly different between conditions. The black dots and dots with black outlines in rhizosphere and root samples indicate ASVs that were also differentially abundant in the nitrate versus native soil. These dots indicate ASVs having a different (black) or similar (black outline) pattern of enrichment/depletion as observed in the soil.

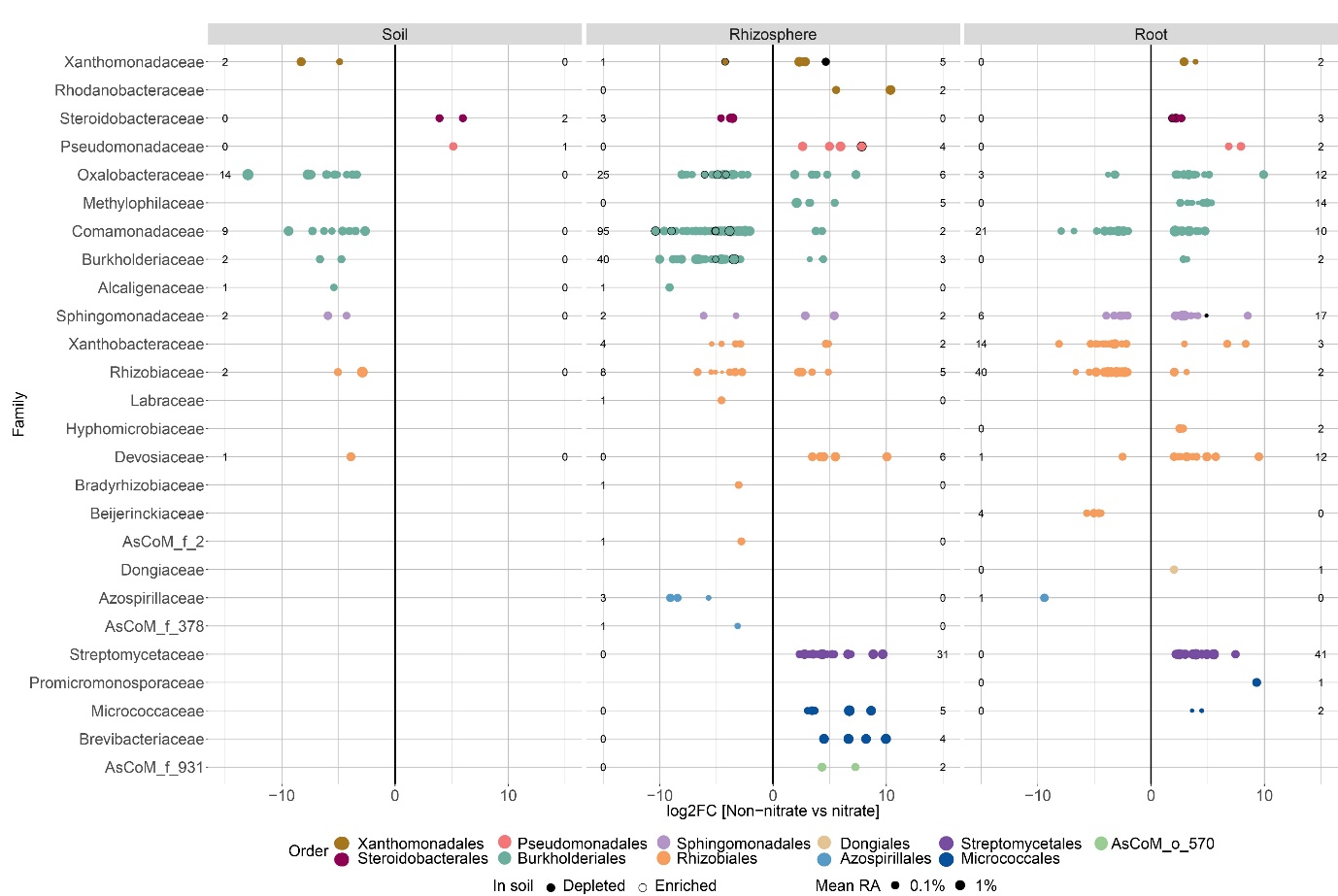

**Supplementary Figure 9. ASVs belonging to different bacterial taxa are differentially enriched in soil, rhizosphere, or roots of *nfr5* plants when grown in native or nitrate supplemented Cologne soil.** Each dot presents an ASV significantly different in abundance in nitrate versus native soil conditions. The size of the dot represents the relative abundance in the condition in which it is enriched. The color of the dots corresponds to the taxonomic order assignment. The numbers at the left/right edge of the plots represent the number of ASVs within the respective family found to be significantly different between conditions. The black dots and dots with black outlines in rhizosphere and root samples indicate ASVs that were also differentially abundant in the nitrate versus native soil. These dots indicate ASVs having a different (black) or similar (black outline) pattern of enrichment/depletion as observed in the soil.

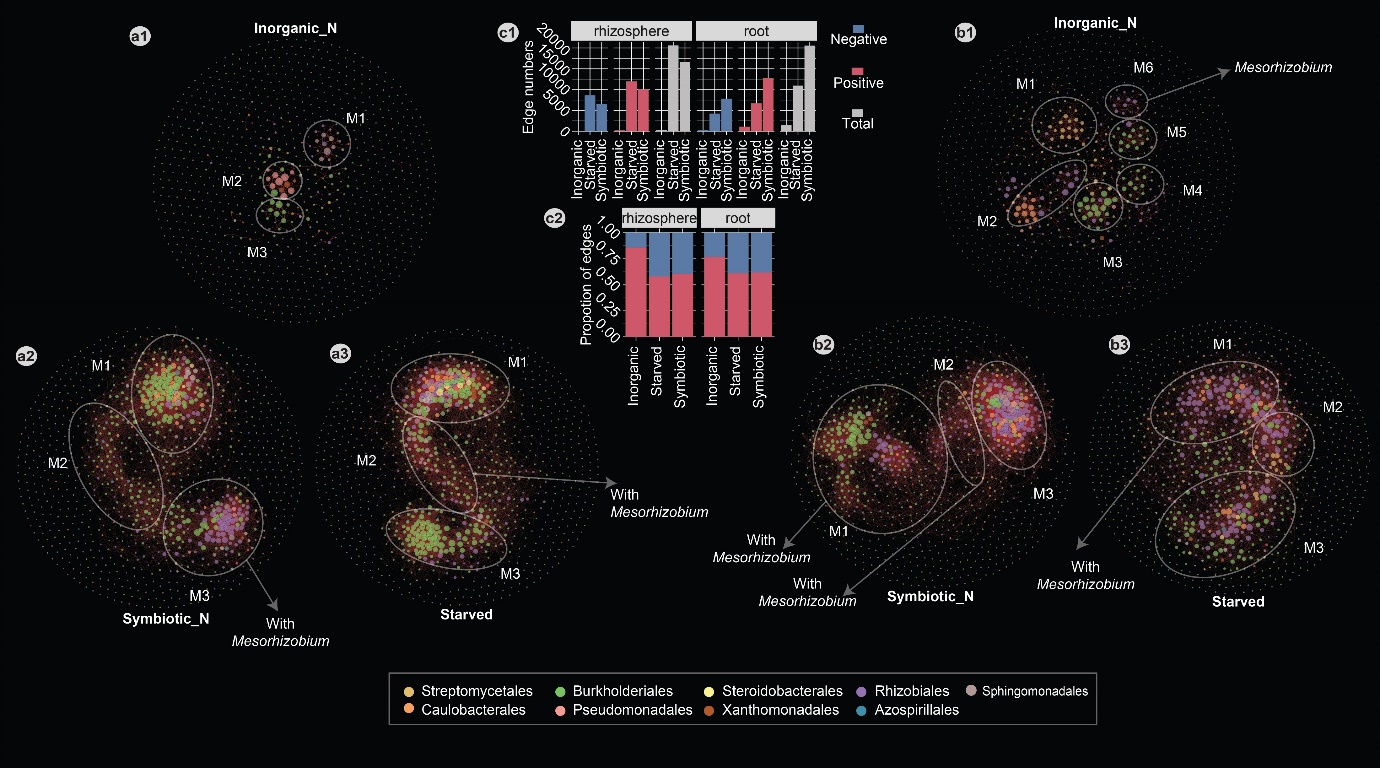

**Supplementary Figure 10. Positive correlation-based networks of ASVs in the rhizosphere (a) and root (b) compartments of plants grown in inorganic nitrogen (a1&b1) or symbiotic nitrogen (a2&b2), as well as starved (a3&b3) conditions**. Each dot represents an ASV. The color of the dots indicates the taxonomic order, size of the dots represents the degree of correlation network. Dotted lines mark clustered modules (M1 to M6). **c)** Number (**c1**) and proportion (**c2**) of positive (red) and negative (blue) edges between ASVs in the three nutritional states.

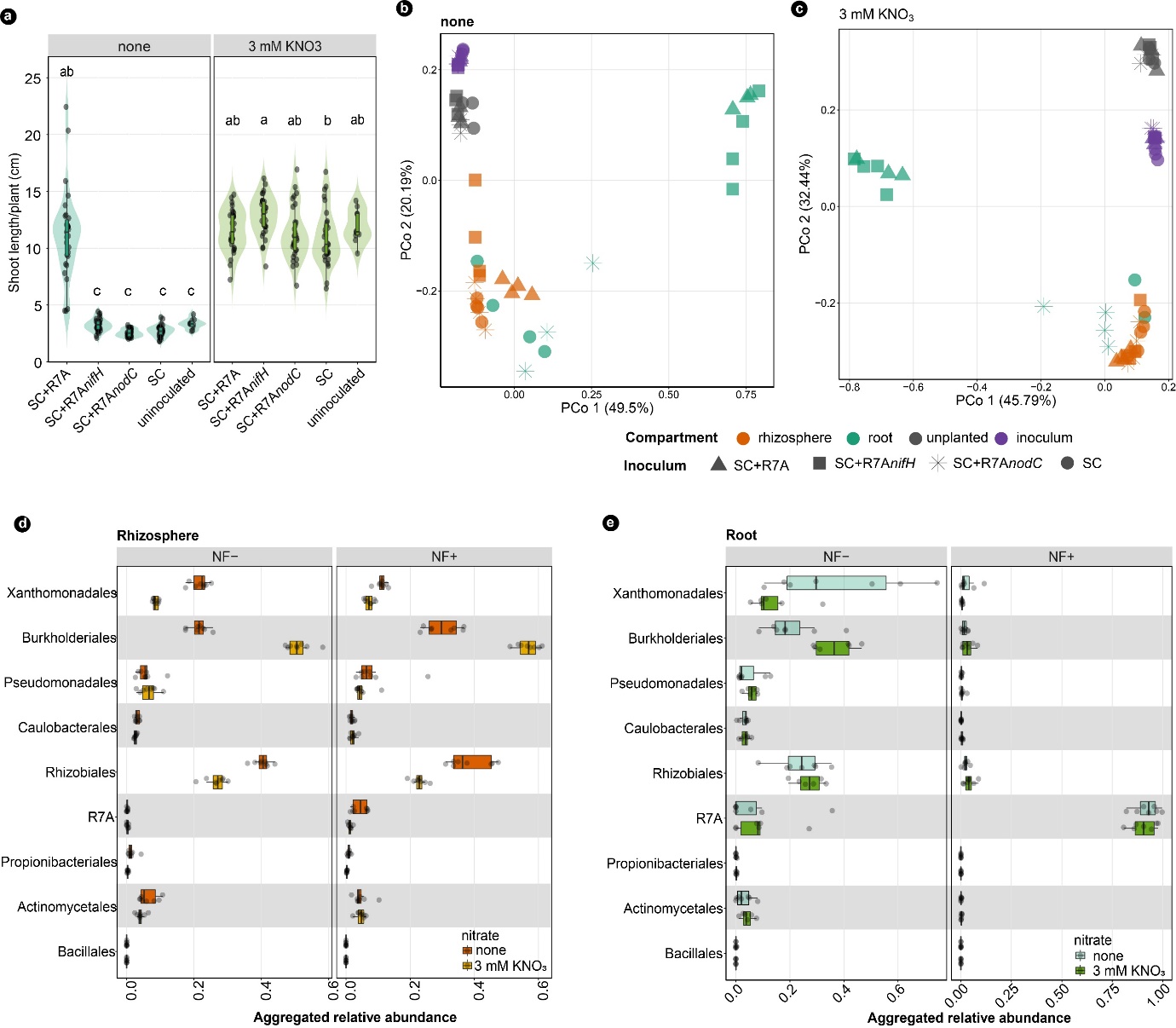

**Supplementary Figure 11. Reconstitution experiments using SynComs reveal that Nod factor production and nitrogen nutrition impact *Lotus* growth and microbiota assembly*.*** (**a**) Shoot length of Gifu grown in different conditions. Letters indicate statistically significant difference (Tukey HSD test, *p*<0.05). PCoA analysis based on Bray-Curtis distances on nitrogen-depleted samples (**b**) and nitrate-supplied samples (**c**). (**d**) Aggregated RA for members of the assigned taxonomic order in the rhizosphere and (**e**) root. NF- indicates samples inoculated with SC+R7A*nodC* and SC, lacking Nod factor production. NF+ indicates samples inoculated with SC+R7A and SC+R7A*nifH* with Nod factor production.

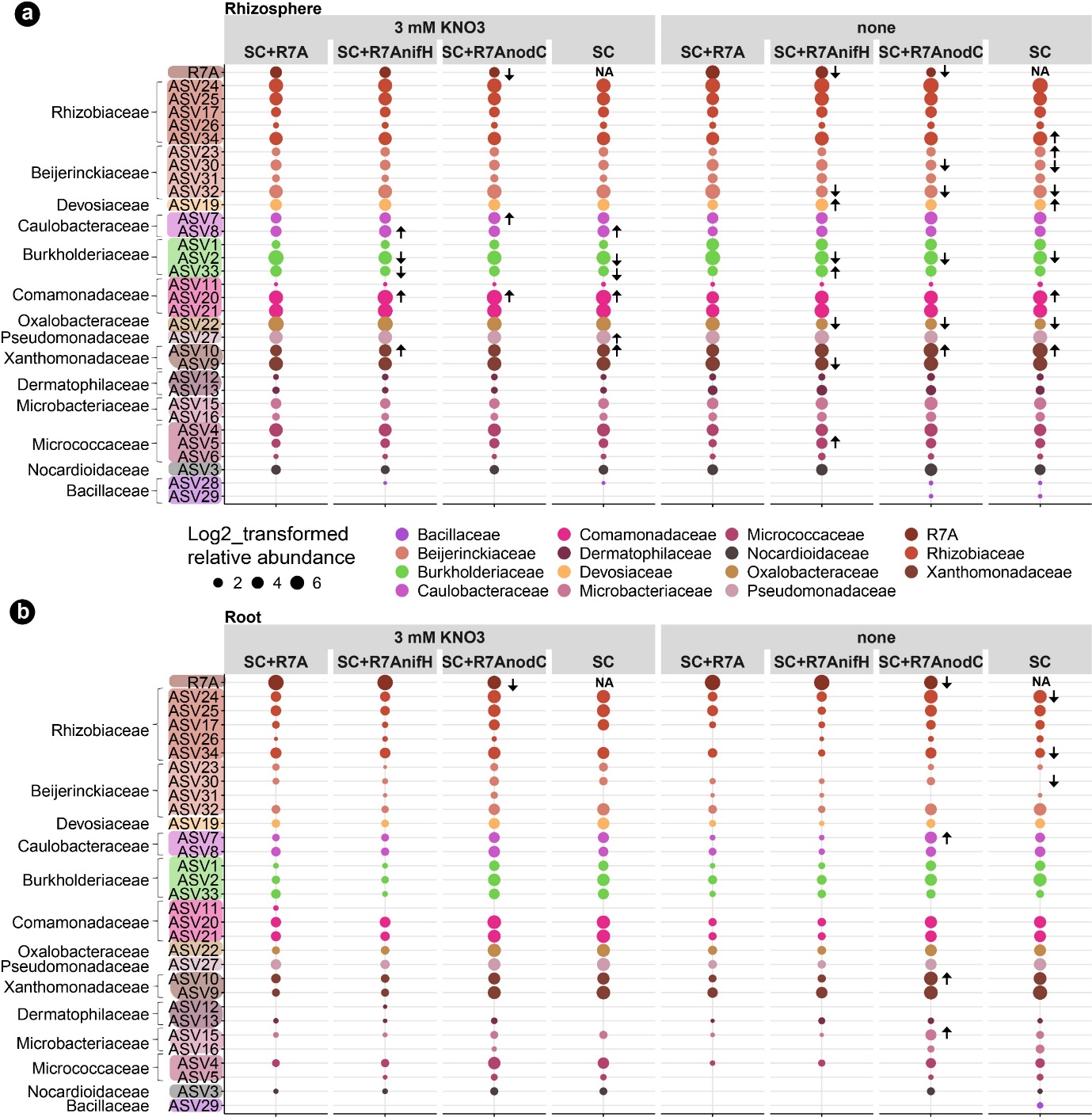

**Supplementary Figure 12. Relative abundance of individual ASVs in reconstitution experiments.** RA of individual ASVs in the **a)** rhizosphere and **b)** root compartments are shown by the size of the dots. The taxonomic assignment (family) is shown by colors. Arrows indicate a significantly higher (upwards) or lower (downwards) RA for the ASVs when part of the indicated inoculum in comparison to its abundance in SC+R7A, and in the respective nutrient condition (none or 3 mM KNO_3_).

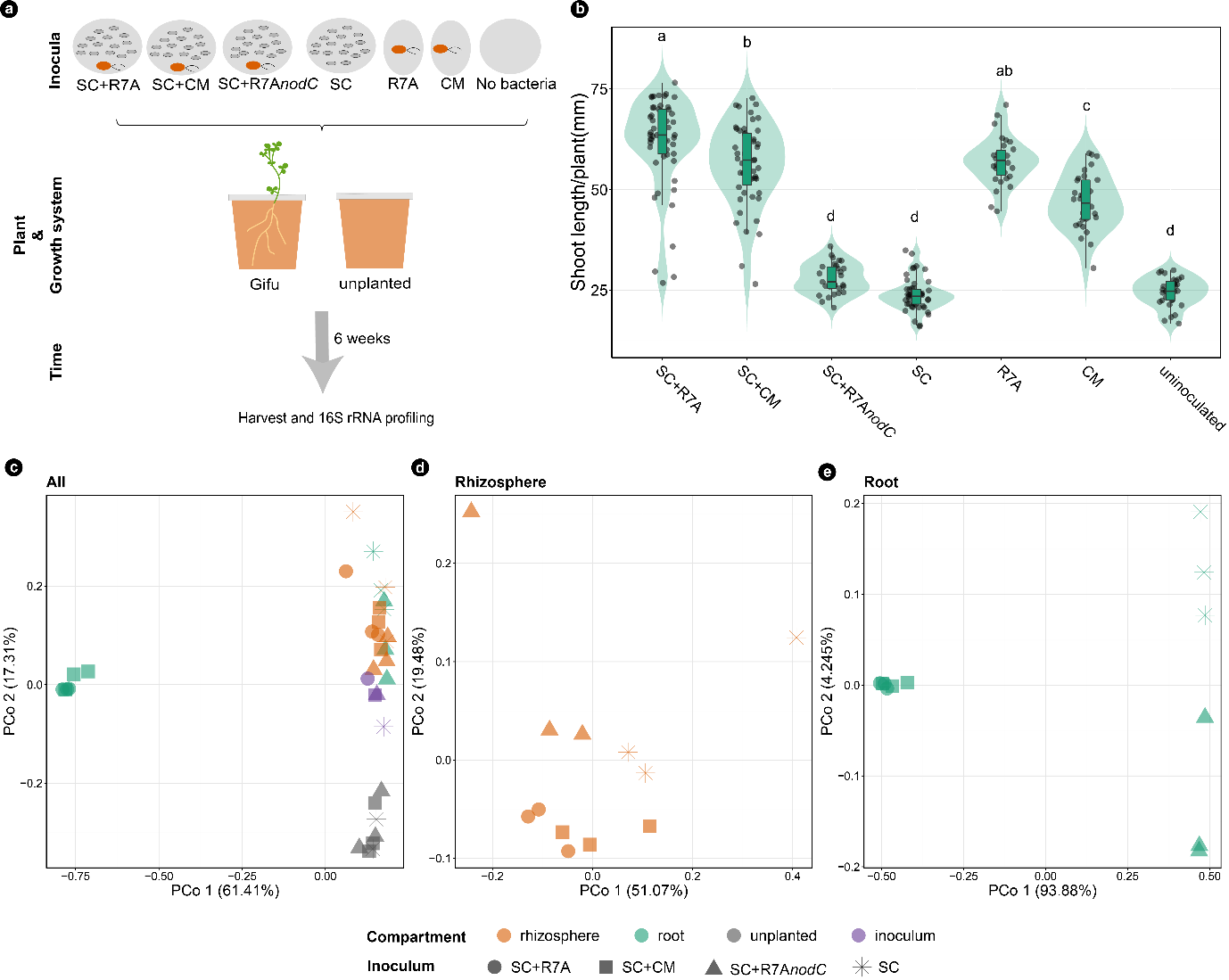

**Supplementary Figure 13. An independent reconstitution experiment using SynComs reveals that bacterial communities of *Lotus* are dependent on Nod factor signaling.** (**a**) Experiment design. (**b**) Shoot length of Gifu grown in different conditions. Letters indicate statistically significant difference (TukeyHSD test, *p*<0.05). PCoA analysis based on Bray-Curtis distances on all samples (**c**), rhizosphere samples (**d**), and root samples (**e**). CM: symbiotic *Mesorhizobium* strain isolated from Cologne soil.

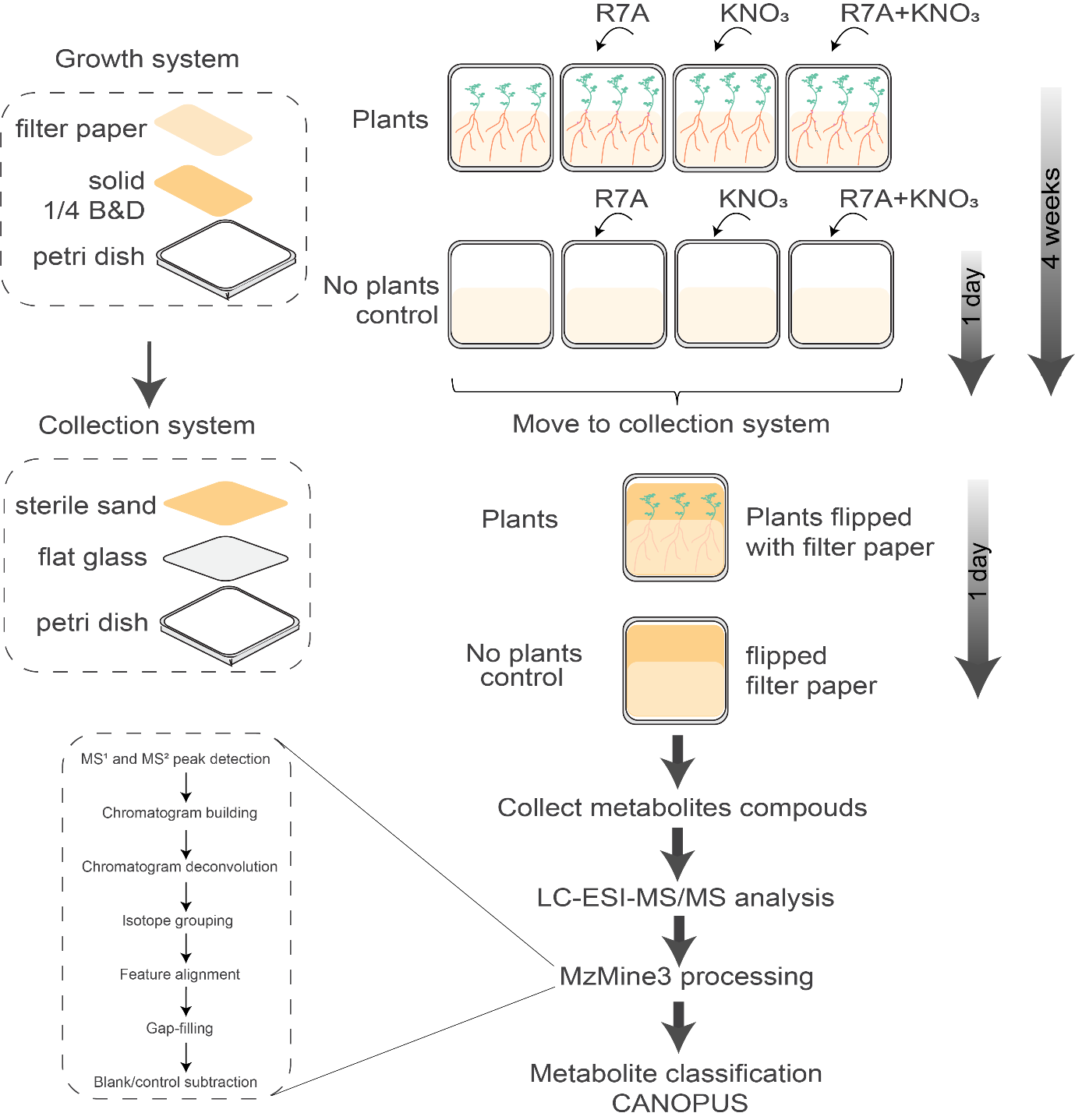

**Supplementary Figure 14. Scheme of the experiment design and process for detecting *Lotus* root exudates.**

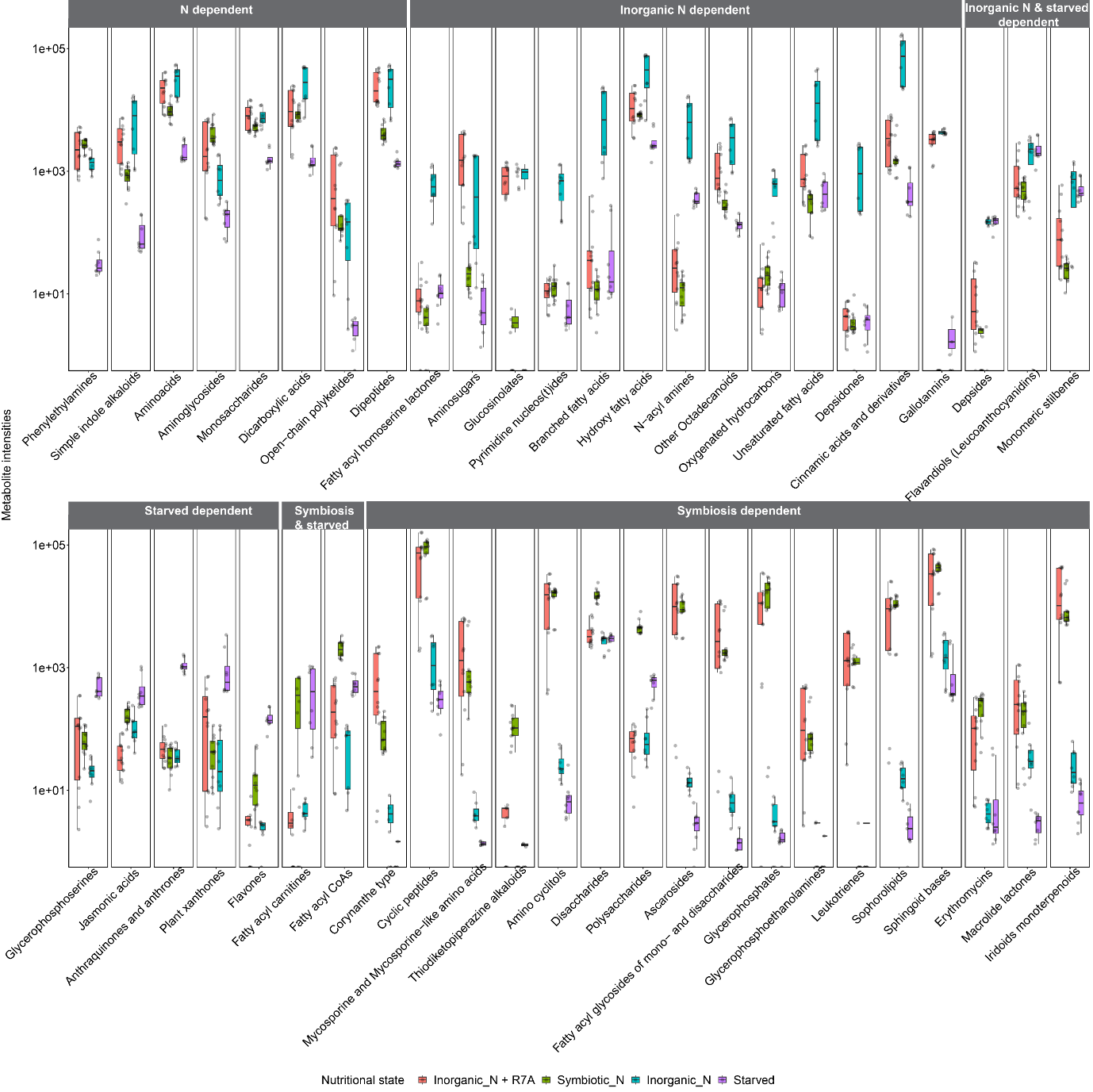

**Supplementary Figure 15. Chemical compound intensity at the most specific class level in the analyzed nutritional states.** The chemical compound identified as “inorganic N dependent”, “N dependent”, “starved dependent”, “symbiosis & starved dependent” and “symbiosis dependent” are shown.
